# Neonatal morphometric similarity mapping for predicting brain age and characterizing neuroanatomic variation associated with preterm birth

**DOI:** 10.1101/569319

**Authors:** Paola Galdi, Manuel Blesa, David Q. Stoye, Gemma Sullivan, Gillian J. Lamb, Alan J. Quigley, Michael J. Thrippleton, Mark E. Bastin, James P. Boardman

## Abstract

Multi-contrast MRI captures information about brain macro- and micro-structure which can be combined in an integrated model to obtain a detailed “fingerprint” of the anatomical properties of an individual’s brain. Inter-regional similarities between features derived from structural and diffusion MRI, including regional volumes, diffusion tensor metrics, neurite orientation dispersion and density imaging measures, can be modelled as morphometric similarity networks (MSNs). Here, individual MSNs were derived from 105 neonates (59 preterm and 46 term) who were scanned between 38 and 45 weeks postmenstrual age (PMA). Inter-regional similarities were used as predictors in a regression model of age at the time of scanning and in a classification model to discriminate between preterm and term infant brains. When tested on unseen data, the regression model predicted PMA at scan with a mean absolute error of 0.70 *±* 0.56 weeks, and the classification model achieved 92% accuracy. We conclude that MSNs predict chronological brain age accurately; and they provide a data-driven approach to identify networks that characterise typical maturation and those that contribute most to neuroanatomic variation associated with preterm birth.

**Highlights:**

1. Multiple MRI features are integrated in a single model to study brain maturation in newborns.
2. Morphometric similarity networks (MSNs) provide a whole-brain description of the structural properties of neonatal brain.
3. The information encoded in MSNs is predictive of chronological brain age in the perinatal period.
4. MSNs provide a novel data-driven method for investigating neuroanatomic variation associated with preterm birth.

## 1. Introduction

Preterm birth is closely associated with increased risk of neurodevelopmental, cognitive and psychiatric impairment that extends across the life course (Nosarti et al., 2012; Anderson, 2014; Mathewson et al., 2017; Van Lieshout et al., 2018). Structural and diffusion MRI (sMRI and dMRI) support the conceptualisation of atypical brain growth after preterm birth as a process characterised by micro-structural alteration of connective pathways due to impaired myelination and neuronal dysmaturation (Boardman et al., 2006; Anjari et al., 2007; Counsell et al., 2008; Ball et al., 2013b; Back and Miller, 2014; Van Den Heuvel et al., 2015; Eaton-Rosen et al., 2015; Thompson et al., 2016; Batalle et al., 2017; Telford et al., 2017; Batalle et al., 2018); this leads to a “dysconnectivity phenotype” that could form the basis for long term functional impairment (Boardman et al., 2010; Caldinelli et al., 2017; Keunen et al., 2017; Cao et al., 2017; Batalle et al., 2018b). However, there has not been a unified approach that incorporates information from sMRI and dMRI to study brain maturation in the perinatal period so the set of image features that best capture brain maturation, and support image classification, are unknown.

The majority of neonatal connectomics studies have used single modes of data such as dMRI tractography (Brown et al., 2014; Batalle et al., 2017; Blesa et al., 2019) or resting-state functional connectivity (Ball et al., 2016; Smyser et al., 2016a). An alternative connectome model is the structural covariance network (SCN) approach (Alexander-Bloch et al., 2013) in which covariance between regional measurements is calculated across subjects, resulting in a single network for the entire population. Other approaches have constructed subject-specific SCNs (Li et al., 2017; Mahjoub et al., 2018) or higher order morphological networks to model the relationship between ROIs across different views (Soussia and Rekik, 2018), but these techniques have been restricted to the use of morphometric variables available through standard structural T1-weighted MRI sequences and by using a single metric (e.g. cortical thickness) to assess the “connectivity” between nodes (Shi et al., 2012).

Based on observations that integrating data from different MRI sequences enhances anatomic characterization (Melbourne et al., 2014; Kulikova et al., 2015; Ball et al., 2017; Thompson et al., 2018a), we investigated whether whole-brain structural connectomes derived from multi-modal data within a prediction framework can capture novel information about perinatal brain development. We used morphometric similarity networks (MSNs) to model inter-regional correlations of multiple macro- and micro-structural multi-contrast MRI variables in a single individual. This approach was originally devised to study how human cortical networks underpin individual differences in psychological functions (Seidlitz et al., 2018), and we adapted it to describe both cortical and subcortical regions in the developing brain. The method works by computing for each region of interest (ROI) a number of metrics derived from different MRI sequences which are arranged in a vector. The aim is to obtain a multidimensional description of the structural properties of the ROIs. The MSN is then built considering the ROIs as nodes and modelling connection strength as the correlation between pairs of ROI vectors, thus integrating in a single connectome the ensemble of imaging features. The pattern of inter-regional correlations can be conceptualised as a “fingerprint” of an individual’s brain.

We investigated the utility of MSNs for describing brain maturation, and for patient classification. The edges of individual MSNs were used to train two predictive models: a regression model to predict postmenstrual age (PMA) at scan and identify the set of image features that best model chronological brain age; and a classification model to discriminate between preterm infants at term equivalent age and term neonates, and thereby identify the networks that explain neuroanatomic variation associated with preterm birth. We hypothesized that predictive models based on MSNs, which integrate information from multiple data modalities, would outperform models based on single metrics and single data modalities.

## 2. Material and methods

### 2.1. Participants and data acquisition

Participants were recruited as part of a longitudinal study designed to investigate the effects of preterm birth on brain structure and long term outcome. The study was conducted according to the principles of the Declaration of Helsinki, and ethical approval was obtained from the UK National Research Ethics Service. Parents provided written informed consent. One hundred and twelve neonates underwent MRI at term equivalent age at the Edinburgh Imaging Facility: Royal Infirmary of Edinburgh, University of Edinburgh, UK, and 105 had multi-modal imaging suitable for MSN analysis (7 acquisitions did not yield usable datasets across all modalities due to motion or wakefulness during one or more sequences). The study group contained 46 term and 59 preterm infants (details are provided in Table 1). The distribution of PMA at scan for all participants, for the term and preterm groups, and the distribution by gender are shown in Fig. 1. Of the preterm infants, 12 had bronchopulmonary dysplasia, 3 had necrotising enterocolitis and 3 required treatment for retinopathy of prematurity.

**Figure 1:**
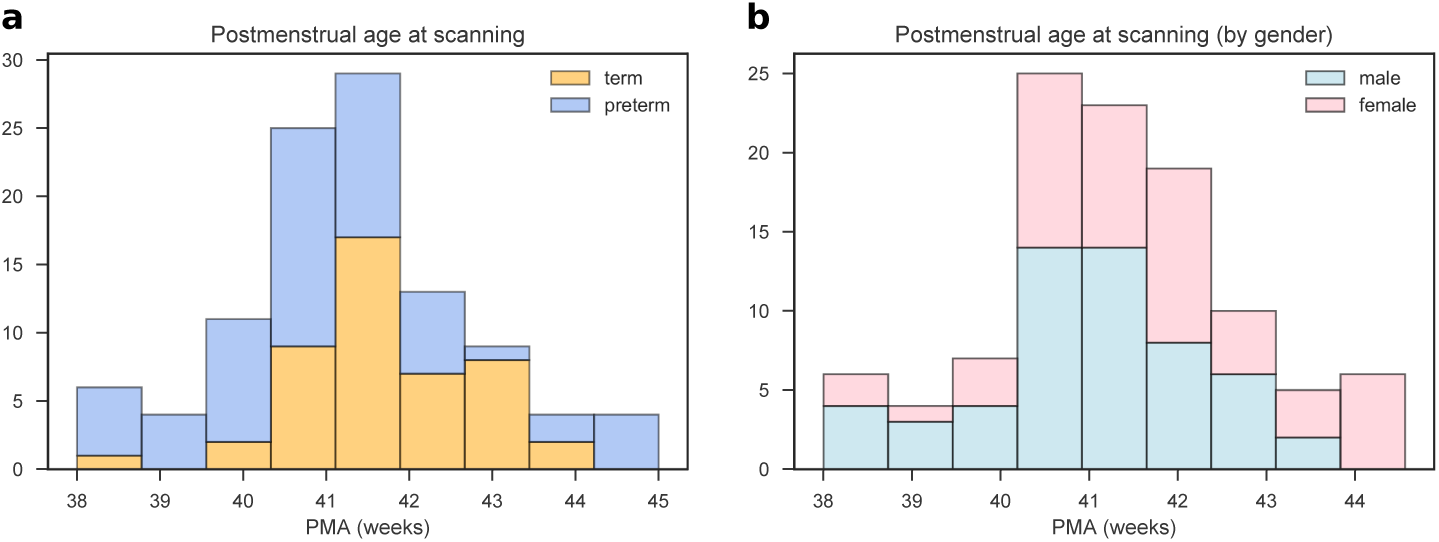
Distribution of postmenstrual age at scan for all subjects. a) Age distribution for the for term (blue) and preterm (orange) groups. b) Age distribution for male (blue) and female (pink) participants.

**Table 1:** Participant characteristics. The last column reports the *p* values of the group differences computed with the Wilcoxon rank-sum test for continuous variables and with the chi-squared test for categorical variables.

|  | preterm (N=59) | term (N=46) | all (N=105) | preterm vs. term |
| --- | --- | --- | --- | --- |
| PMA at birth (weeks) | 23.42-32.00 | 37.00-42.00 | 23.42-42.00 | $p = 1.88 \times 10^{-18}$ |
| Birth weight (grams) | 454-2100 | 2556-4560 | 454-4560 | $p = 1.93 \times 10^{-18}$ |
| PMA at scan (weeks) | 38.00-44.56 | 38.28-43.84 | 38.00-44.56 | $p = 0.0035$ |
| M:F ratio | 29:30 | 26:20 | 55:50 | $p = 0.4532$ |

PMA = Postmenstrual age, M = male, F = female.

A Siemens MAGNETOM Prisma 3 T MRI clinical scanner (Siemens Healthcare Erlangen, Germany) and 16-channel phased-array paediatric head coil were used to acquire: 3D T1-weighted MPRAGE (T1w) (acquired voxel size = 1mm isotropic) with TI 1100 ms, TE 4.69 ms and TR 1970 ms; 3D T2-weighted SPACE (T2w) (voxel size = 1mm isotropic) with TE 409 ms and TR 3200 ms; and axial dMRI. dMRI was acquired in two separate acquisitions to reduce the time needed to re-acquire any data lost to motion artefact: the first acquisition consisted of 8 baseline volumes (b = 0 s/mm^2^ [b0]) and 64 volumes with b = 750 s/mm^2^, the second consisted of 8 b0, 3 volumes with b = 200 s/mm^2^, 6 volumes with b = 500 s/mm^2^ and 64 volumes with b = 2500 s/mm^2^; an optimal angular coverage for the sampling scheme was applied (Caruyer et al., 2013). In addition, an acquisition of 3 b0 volumes with an inverse phase encoding direction was performed. All dMRI images were acquired using single-shot spin-echo echo planar imaging (EPI) with 2-fold simultaneous multislice and 2-fold in-plane parallel imaging acceleration and 2 mm isotropic voxels; all three diffusion acquisitions had the same parameters (TR/TE 3400/78.0 ms).

Infants were fed and wrapped and allowed to sleep naturally in the scanner. Feeds were timed to increase the likelihood of post-prandial sleep, flexible earplugs and neonatal earmuffs (MiniMuffs, Natus) were used for acoustic protection, and a soothing environment was created in terms of light and noise. Pulse oximetry, electrocardiography and temperature were monitored. All scans were supervised by a doctor or nurse trained in neonatal resuscitation. Each acquisition was inspected contemporaneously for motion artefact and repeated if there had been movement but the baby was still sleeping; dMRI acquisitions were repeated if signal loss was seen in 3 or more volumes. The majority of the cohort had one or more sequences repeated in order to acquire the best possible quality data for processing.

Conventional images were reported by an experienced paediatric radiologist (A.J.Q.) using a structured system (Leuchter et al., 2014; Woodward et al., 2006), and none of the images included in the final sample (*N* = 105) showed evidence of focal parenchymal injury (defined as post-haemorrhagic ventricular dilatation, porencephalic cyst or cystic periventricular leukomalacia), or central nervous system malformation.

### 2.2. Data preprocessing

All the following preprocessing steps, including maps calculation and quality check, were performed using dcm2niix, FSL, MRtrix, MIRTK, ANTs, Connectome Workbench and cuDIMOT (Smith et al., 2004; Avants et al., 2011; Marcus et al., 2011; Makropoulos et al., 2014; Li et al., 2016; Hernandez-Fernandez et al., 2019; Tournier et al., 2019).

First, all DICOM image files (dMRI and sMRI) were converted to NIFTI (Li et al., 2016). Structural data were preprocessed using the developing Human Connectome Project (dHCP) minimal structural processing pipeline for neonatal data (Makropoulos et al., 2018). Briefly, the T1w image was co-registered to the T2w image, both were corrected for bias field inhomogeinities (Tustison et al., 2010) and an initial brain mask was created (Smith, 2002). Following this, the brain was segmented into different tissue types (CSF: cerebrospinal fluid; WM: white matter; cGM: cortical grey matter; GM: subcortical grey matter) using the Draw-EM algorithm (Makropoulos et al., 2014). Twenty manually labelled atlases (Gousias et al., 2012) were then registered to each subject using a multi-channel registration approach, where the different channels of the registration were the original intensity T2-weighted images and GM probability maps. These GM probability maps were derived from an initial tissue segmentation, performed using tissue priors propagated through registration of a preterm probabilistic tissue atlas (Serag et al., 2012). The framework produces several output files, but for this study only the aligned T1w and the T2w images and the parcellation in 87 ROIs were used (Makropoulos et al., 2018). Note that from these 87 ROIs six were removed: the background, the unlabelled brain area (mainly internal capsule), the CSF, the lateral ventricles (left and right) and the corpus callosum (see section 2.4).

Diffusion MRI processing was performed as follows: for each subject the two dMRI acquisitions were first concatenated and then denoised using a Marchenko-Pastur-PCA-based algorithm (Veraart et al., 2016, b); the eddy current, head movement and EPI geometric distortions were corrected using outlier replacement and slice-to-volume registration with TOPUP and EDDY (Andersson et al., 2003; Smith et al., 2004; Andersson and Sotiropoulos, 2016; Andersson et al., 2016, 2017); bias field inhomogeneity correction was performed by calculating the bias field of the mean b0 volume and applying the correction to all the volumes (Tustison et al., 2010). This framework only differs from the optimal pipeline for diffusion preprocessing presented in Maximov et al. (2019) in that we did not perform the final smoothing or the gibbs-ring removal (Kellner et al., 2016) due to the nature of the data (partial fourier space acquisition).

The mean b0 EPI volume of each subject was co-registered to their structural T2w volume using boundary-based registration (Greve and Fischl, 2009), then the inverse transformation was used to propagate ROI labels to dMRI space, with a modified bbrslope parameter of 0.5, which is used for neonatal data (Toulmin et al., 2015).

For each ROI, two metrics were computed in structural space: ROI volume and the mean T1w/T2w signal ratio (Glasser and Van Essen, 2011). The other ten metrics were calculated in native diffusion space: five metrics derived from the diffusion kurtosis (DK) model (Jensen et al., 2005) and five derived from the Neurite Orientation Dispersion and Density Imaging model (NODDI) (Zhang et al., 2012; Tariq et al., 2016).

### 2.3. Feature extraction

#### 2.3.1. Structural metrics

ROI volumes were calculated without normalising for the whole brain volume, as they are used only to compute inter-regional similarities within subjects. The mean T1w/T2w signal ratio was calculated before the bias field correction. The T1w/T2w ratio was used because it enhances myelin contrast and mathematically cancels the signal intensity bias related to the sensitivity profile of radio frequency receiver coils (Glasser and Van Essen, 2011).

#### 2.3.2. Diffusion kurtosis metrics

The diffusion kurtosis (DK) model is an expansion of the diffusion tensor model. In addition to the diffusion tensor, the DK model quantifies the degree to which water diffusion in biological tissues is non-Gaussian using the kurtosis tensor. The reason for this is that the Gaussian displacement assumption underlying the diffusion tensor breaks at high b-values (Jensen et al., 2005). On the kurtosis component, we only focus on the mean value along all diffusion directions.

The metrics obtained from the DK model for each ROI are the means of: the fractional anisotropy (FA), mean, axial and radial diffusivity (MD, RD, AD) and kurtosis (MK). The MK map quantifies the deviation from Gaussianity of water molecule displacement and can reflect different degrees of tissue heterogeneity (Steven et al., 2014).

#### 2.3.3. NODDI metrics

We included NODDI metrics alongside the more commonly adopted diffusion tensor measures as previous studies have shown that NODDI indices are sensitive to underlying biological changes in the brain and provide more specific microstructural characteristics, in agreement with histology (Grussu et al., 2017; Batalle et al., 2018).

For the NODDI measures, the Bingham distribution was employed (Tariq et al., 2016) as it allows extra flexibility by describing fibre dispersion along two orthogonal axes. From this NODDI implementation we obtain five metrics: intracellular volume fraction (*υ*_ic_), isotropic volume fraction (*υ*_iso_), the orientation dispersion index along the primary and secondary directions (ODI_P_ and ODI_S_) and the overall orientation dispersion index (ODI_TOT_).

One limitation of this model is that it requires fixing a value for the diffusivity along the axons. However, optimal values for this parameter are region-dependent (Karmacharya et al., 2018) and the default value may be suboptimal for the neonatal population as it has been optimised using an adult cohort (Zhang et al., 2012; Karmacharya et al., 2018). Several studies have been reporting NODDI values for neonates using default (or unspecified) parameters (Batalle et al., 2018; Bastiani et al., 2018; Karmacharya et al., 2018) or modified ones (Kunz et al., 2014; Jelescu et al., 2015). As our goal was not to report NODDI values for the different areas, and because of the lack of reference values for this population, we calculated NODDI maps using default parameters (Batalle et al., 2018).

### 2.4. Data Quality Control

The parcellations obtained after the processing were visually inspected and parcels corresponding to CSF and background parcels were excluded because they do not represent brain tissue. We observed a poor segmentation of the corpus callosum in part of the subjects, but we did not find any anomalies in the rest of the parcels. This effect could be caused by different factors: a) this area is problematic to segment due to the proximity to CSF and its small thickness (see for example Otsuka et al. (2019)); b) the framework we used was optimised for the dHCP data that have a very high resolution (0.5 mm^3^ isotropic) and data quality, making the partial volume effect more noticeable in data with a resolution of 1 mm^3^; c) or susceptibility artifacts. Instead of removing the subjects with a poor segmentation, we decided to remove the corpus callosum from the model, aiming at maximising the number of subjects. As a result of the whole quality check, we include the whole population (*N* = 105) and each network is composed of 81 nodes (ROIs).

For the dMRI data we use eddy QC (Bastiani et al., 2019). The quality control is performed at subject level and group level. Eddy QC provides several measures related to the rotation, translation and outliers of the images. In addition, it also computes the signal-to-noise (SNR) ratio maps of the b0 volumes and the contrast-to-noise (CNR) ratio maps for the different b-values. These maps can be used at group level to visualise the quality of the data (Bastiani et al., 2018). The results show that the overall quality of the data-set was good (Fig. 2). For eddy QC to work, we removed the b-value = 200 s/mm^2^ only from the quality control. This is because the low number of volumes with this b-value sometimes leads the Gaussian process performed by eddy to produce a perfect fit, which makes the CNR maps unrealistic.

**Figure 2:**
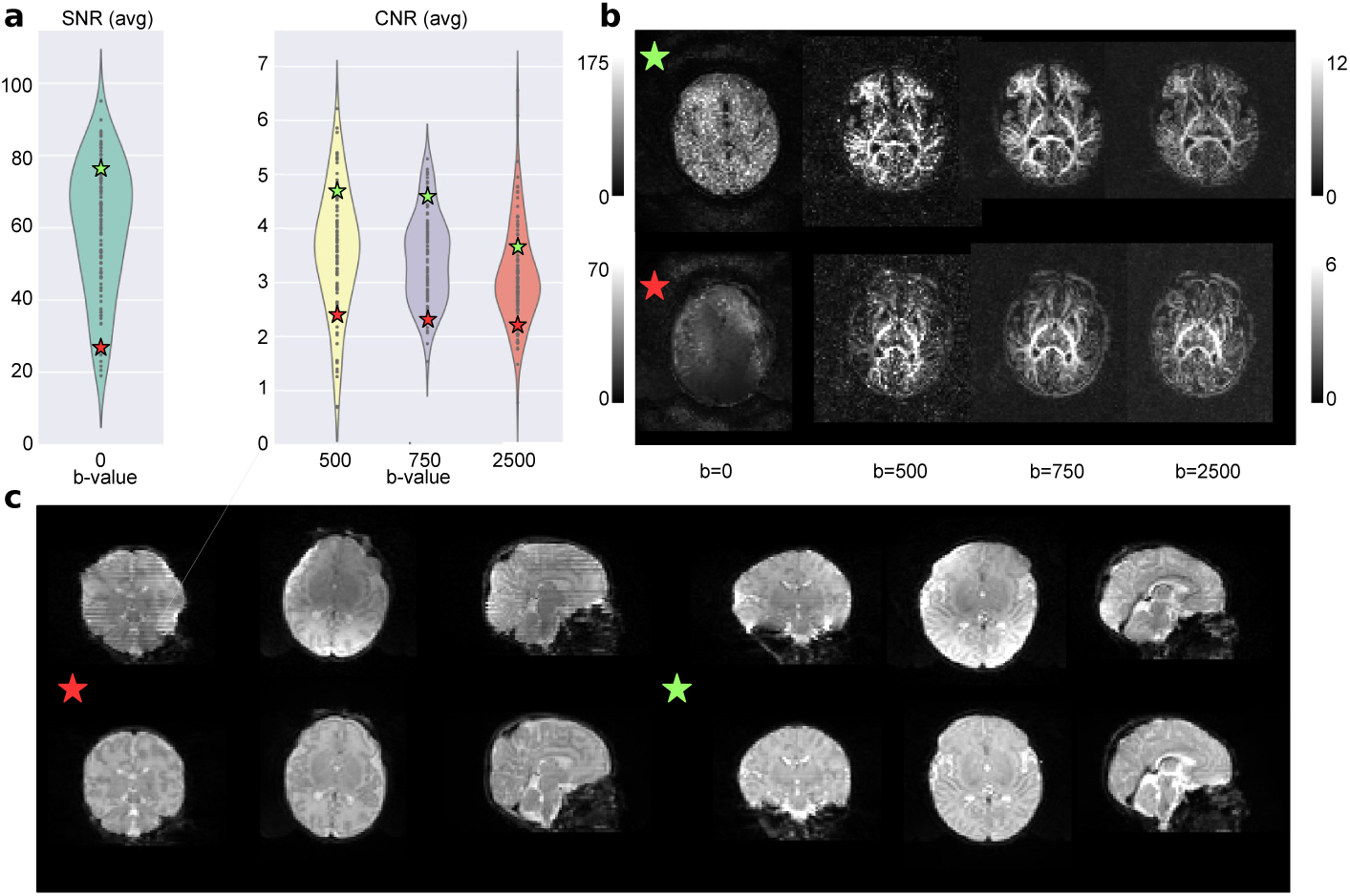
Quality control results. a) Results for the overall population with two selected subjects, one from the top quartile of the SNR and CNR distributions (green star) and the other from the bottom quartile (red star). b) The SNR and CNR maps for the selected subjects. c) The b0 of both subjects before and after the processing pipeline.

Fig. 2 shows two representative subjects, one from the top quartile of the SNR and CNR distributions (green star) and one from the bottom quartile (red star). In the first panel we can see where they are placed in terms of SNR and CNR over the overall population. The second panel shows the SNR maps (for the b0) and the CNR maps (for the rest of b-values). The bottom panel of the Fig. 2 shows the b0 before and after the processing of the selected subjects. It is possible to observe the effect of the different steps involved, such as the EPI geometric corrections or the bias field inhomogeneity correction. Supplementary Figs. S8 and S9 report the above results for the term and preterm population respectively.

Following Bastiani et al. (2019), for each volume, motion is quantified by averaging voxel displacement across all voxels (computed as 3 translations and 3 rotations around the x, y and z axes). Absolute displacement is computed with respect to the reference volume, while relative displacement is computed with respect to the previous volume. A summary measure for each subject is calculated as the average (absolute or relative) displacement across all volumes. In Supplementary Fig. S10 we show the distribution of absolute and relative motion for the term and the preterm groups. We compared the distributions with a Wilcoxon rank-sum test and found no difference between the relative motion scores (*W* = 1330*, p* = 0.43) and a significant difference between the absolute motion scores (*W* = 1720*, p* = 0.02). However, as the violin plot shows, this difference is driven by the presence of outliers.

### 2.5. Experimental design and statistical analysis

The models and the analyses described in this section were implemented in Python (v3.6.4) using open source libraries and frameworks for scientific computing, including SciPy (v1.0.0), Numpy (v1.14.0), Statsmodels (v0.8.0), Pandas (v0.22.0), Scikit-learn (v0.19.1) and Matplotlib (v2.1.2) (Jones et al., 2001; Hunter, 2007; Seabold and Perktold, 2010; McKinney et al., 2010; Pedregosa et al., 2011; Van Der Walt et al., 2011).

#### 2.5.1. Network Construction

The MSN for each subject was constructed starting from 81 ROIs; each of the ROI metrics was normalised (z-scored) and Pearson correlations were computed between the vectors of metrics from each pair of ROIs. In this way, the nodes of each network are the ROIs and the edges represent the morphometric similarity between the two related ROIs (Fig. 3). In the following, the terms “edge”, “connection” and “inter-regional similarity” are used interchangeably to refer to the correlation between the regional metrics of a pair of ROIs.

**Figure 3:**
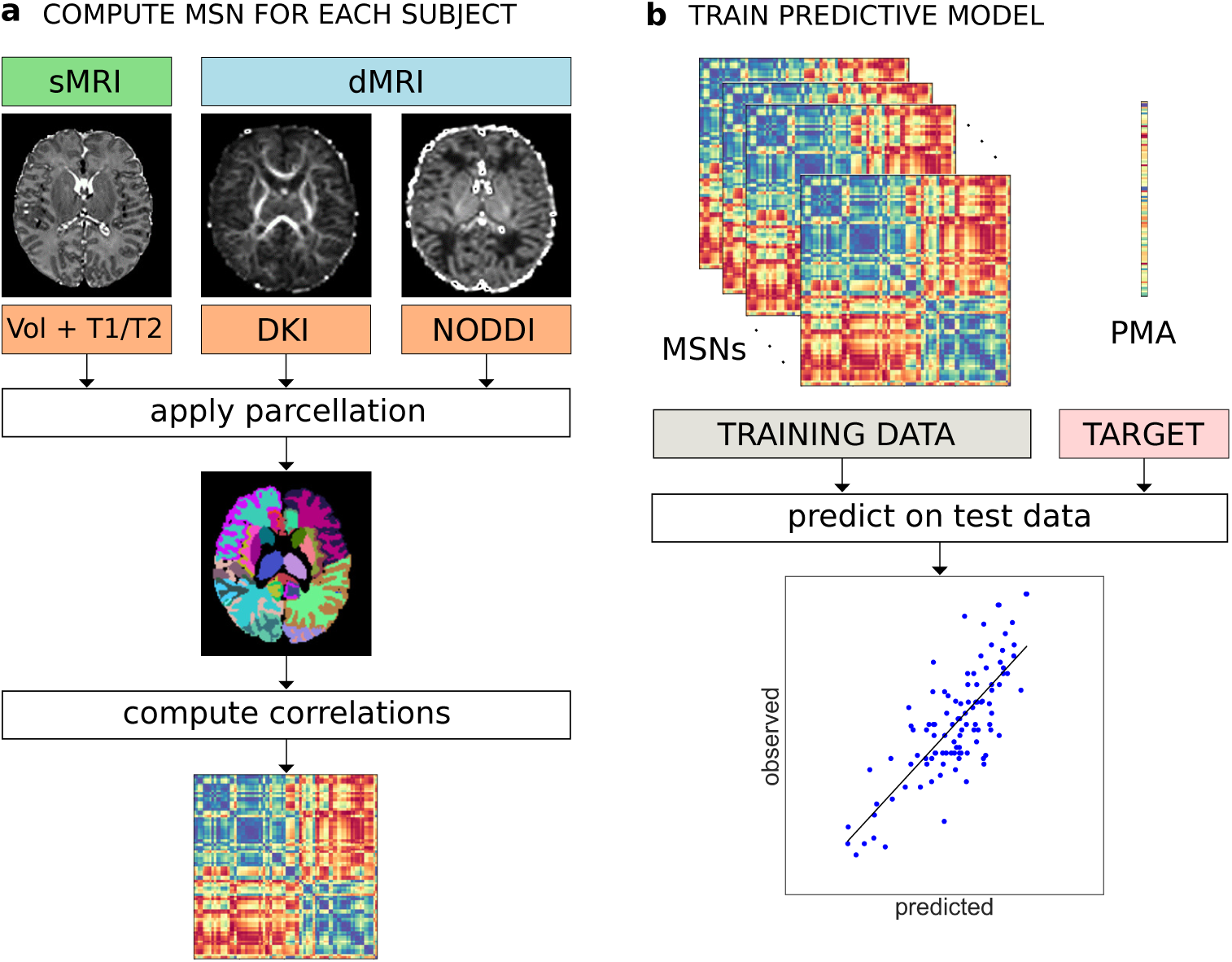
a) Individual MSN construction. Different metrics are extracted from dMRI and sMRI data. The same parcellation is applied to all image types and the average metric values are computed for each ROI. A MSN (represented here as a connectivity matrix) is built by computing the Pearson correlation between the vectors of metrics of each pair of ROIs. b) Training of a predictive model (here for PMA at scan) from individual MSNs. The inter-regional correlations are used as predictor variables in a machine learning model. The performance of the model is evaluated on an independent test set.

#### 2.5.2. Confounding variables

Early exposure to the extrauterine environment due to preterm birth exposes infants to several processes that are known to impact brain maturation (e.g. specific co-morbidities such as bronchopulmonary dysplasia and necrotising enterocolitis (Barnett et al., 2018)), and other processes and diseases that can modify brain maturation (for example gestational age at birth, chorioamnionitis, fetal growth restriction, nutritional insufficiency, pain and medication exposures (Duerden et al., 2016; Anblagan et al., 2016; Barnett et al., 2018; Schneider et al., 2018; Duerden et al., 2018; Blesa et al., 2019)). In addition, there may be as yet unknown environmental risks to brain structural connectivity and genomic and epigenomic factors may interact with gestational age at birth to confer risk (Batalle et al., 2017, 2018b; Boardman et al., 2014; Sparrow et al., 2016; Krishnan et al., 2017). Therefore, it is not possible to define a preterm infant cohort without any exposures to processes that could influence brain maturation. As our intention was to develop an integrated approach for characterising dysmaturation in a study group representative of the target population, rather than to investigate possible drivers of dysmaturation, we did not control for any of the above factors.

We did however find that the preterm group was characterised by higher inscanner motion than the term-group, hence we considered absolute displacement as a confounder (section 2.4). We also observed a positive correlation (*ρ* = 0.27*, p* = 0.0048) between PMA at scan and PMA at birth and a negative correlation (*ρ* = *−*0.22*, p* = 0.0233) between PMA at scan and gender (coded as a binary variable where 0 indicates female infants and 1 male infants), implying that in our sample term subjects and female subjects tend to have their scan acquired at a later age (see also fig. 1). To control for potential bias, we used these confounders as predictors and compared their predictive performance with our network-based features. We tested the interaction between gender and prematurity in a linear regression model of PMA at scan, but the interaction term was not significant (*p* = 0.9634). Birthweight was not included explicitly as a confounder due to its collinearity with PMA at birth.

#### 2.5.3. Regression model for age

We trained a linear regression model with elastic net regularisation to predict PMA at scan – i.e. chronological brain age – in both preterm and term infants starting from individual MSNs. This model was chosen for its ability to cope with a high number of features (Zou and Hastie, 2005). For each subject, the edges of the MSN (inter-regional correlations) were concatenated to form a feature vector to be given as input to the regression model. Since the connectivity matrix representing the MSN is symmetric, we considered only the upper triangular matrix for each subject. Gender and age at birth were included in the model to control for their possible confounding effects. The prediction performances were evaluated with a leave-one-out cross-validation (LOOCV) scheme, by computing the mean absolute error (MAE) averaged across subjects. Within each fold of the LOOCV, the parameters of the elastic net were selected with a nested 3-fold cross-validation loop; the folds were stratified in percentiles to include samples covering the whole age range in each of the folds. Permutation testing was used for the statistical validation of the model performance: the null distribution was built by running the age prediction analysis on 1000 random permutation of the PMA.

#### 2.5.4. Classification model

A Support Vector Machine (SVM) classifier with linear kernel was trained to discriminate between preterm and term infants. As per the regression model, the input for each subject consisted of inter-regional connections taken from the upper triangular connectivity matrix and the performances were evaluated with LOOCV. Age at the time of scanning, gender and motion were included as additional covariates.While in the case of regression the elastic net regularisation performs automatically a variable selection step, recursive feature elimination (RFE) was applied in combination with SVM to select the best subset of connections. Model selection was implemented using nested cross validation: an outer 3-fold cross-validation loop was used to select the SVM parameters and an inner 4-fold cross-validation loop was used for RFE. Folds were stratified to include the same proportion of term and preterm subjects. The accuracy of the model was computed as the number of correctly classified subjects across the leave-one-out folds over the total number of subjects in the test set. The null distribution was built by repeating the exact same analysis 1000 times after randomly assigning subjects to the term and the preterm group.

#### 2.5.5. Feature selection

After the preprocessing phase, twelve different metrics were available for each ROI. To study which combination of features produced better performance in the prediction tasks, we implemented a sequential backward-forward feature selection scheme. Starting from the full set of features, at each iteration we compare the performances of different models built by removing in turn each of the features from the current set of candidate features. We then exclude from the next iteration the feature whose subtraction caused the least increase in prediction error (down to three features, for a total of 73 combinations). The rationale behind this scheme is to explore the space of possible models without enumerating all possible solutions, thus reducing the computational demands compared to an exhaustive search. The procedure was performed separately for the regression and the classification models.

#### 2.5.6. Cross-validation strategy

We adopted LOOCV to select the best performing model in both the age prediction and the classification tasks as this scheme enabled maximum size of the training set and therefore best use of available data, but this strategy might induce high variance in the estimation of prediction accuracy (Kohavi, 1995; Efron, 1983). In the context of brain decoding (i.e. predictions from brain images or signals), LOOCV was shown to produce overly optimistic estimates of prediction accuracy in the within-subject setting (i.e. when all samples are highly correlated because they come from the same subject). In the between-subject setting (as in this work), the performance of LOOCV is similar to schemes involving random splits and mostly determined by sample size (Varoquaux et al., 2017; Varoquaux, 2018). To assess the stability of our results with respect to the chosen cross-validation scheme, we report the prediction accuracy computed with a 10 repeated stratified 5-fold scheme (10-5-fold) for all the models selected with LOOCV.

#### 2.5.7. Comparison with individual metrics and single data modalities models

We compared the performances of the best performing models based on MSNs with three classes of baseline models: a) models based on single global brain metrics (total brain volume and median FA in the WM); b) models based on individual metrics, where instead of similarities, predictors are the concatenation of all regional values for each of the individual metrics used to build MSNs; c) single data modality MSNs, i.e. models built on structural features only (Volume and T1/T2), on DKI features only, and on NODDI features only.

### 2.6. Data and code availability

Source code implementing the methods described in this paper is available upon request to the corresponding author. The preprocessed and anonymised data used in the analyses can be requested through the Brains Image Bank (https://www.brainsimagebank.ac.uk/) (Job et al., 2017).

## 3. Results

### 3.1. Feature selection

In Fig. 4 we report two histograms summarising the LOOCV performance of the 73 different models compared per each task in the backward feature selection scheme. In both cases, we can observe that the models based on all three data modalities achieved better results in terms of prediction accuracy. The performances of each of the compared model are reported in Supplementary Figs. S1 and S3 for the age prediction and for the classification models, respectively.

**Figure 4:**
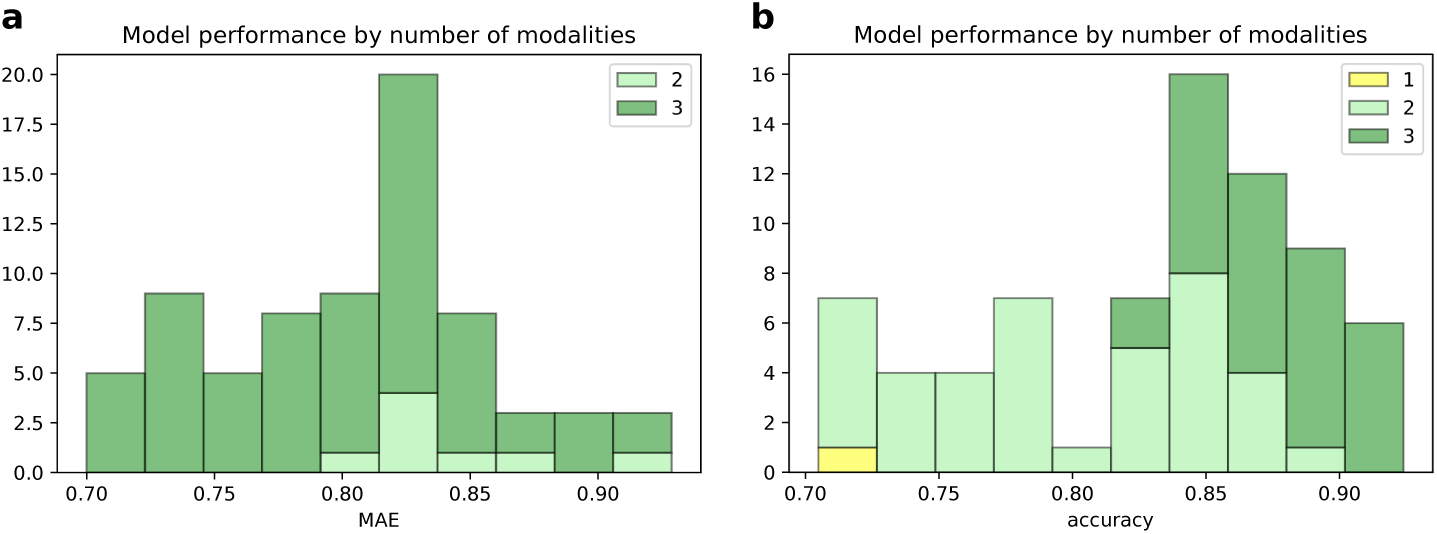
Histograms of the performance of the 73 models compared in the backward feature selection scheme for the age prediction task (a) and for the classification task (b). Bars are grouped by the number of modalities included in the models.

The best performing model for age prediction, which was adopted for all subsequent analyses, was based on seven features (Volume, FA, MD, AD, MK, *υ*_iso_, ODI_P_). Fig. 5a shows the average MSN matrix computed across all subjects for the selected set of features and the matrix of correlation between inter-regional similarities and PMA at scan across subjects. The average MSN matrix shows four main blocks that correspond roughly to positive correlations between ROIs within GM and between ROIs within WM, and to negative correlation between WM ROIs and GM ROIs, indicating that ROIs within GM (and within WM) share similar structural properties, while GM and WM regional descriptors tend to be anti-correlated. The four-block structure is recognisable also in the matrix reporting correlations with chronological age: with increasing age regions within GM or within WM become more similar with each other, while the dissimilarities between GM and WM ROIs increases.

**Figure 5:**
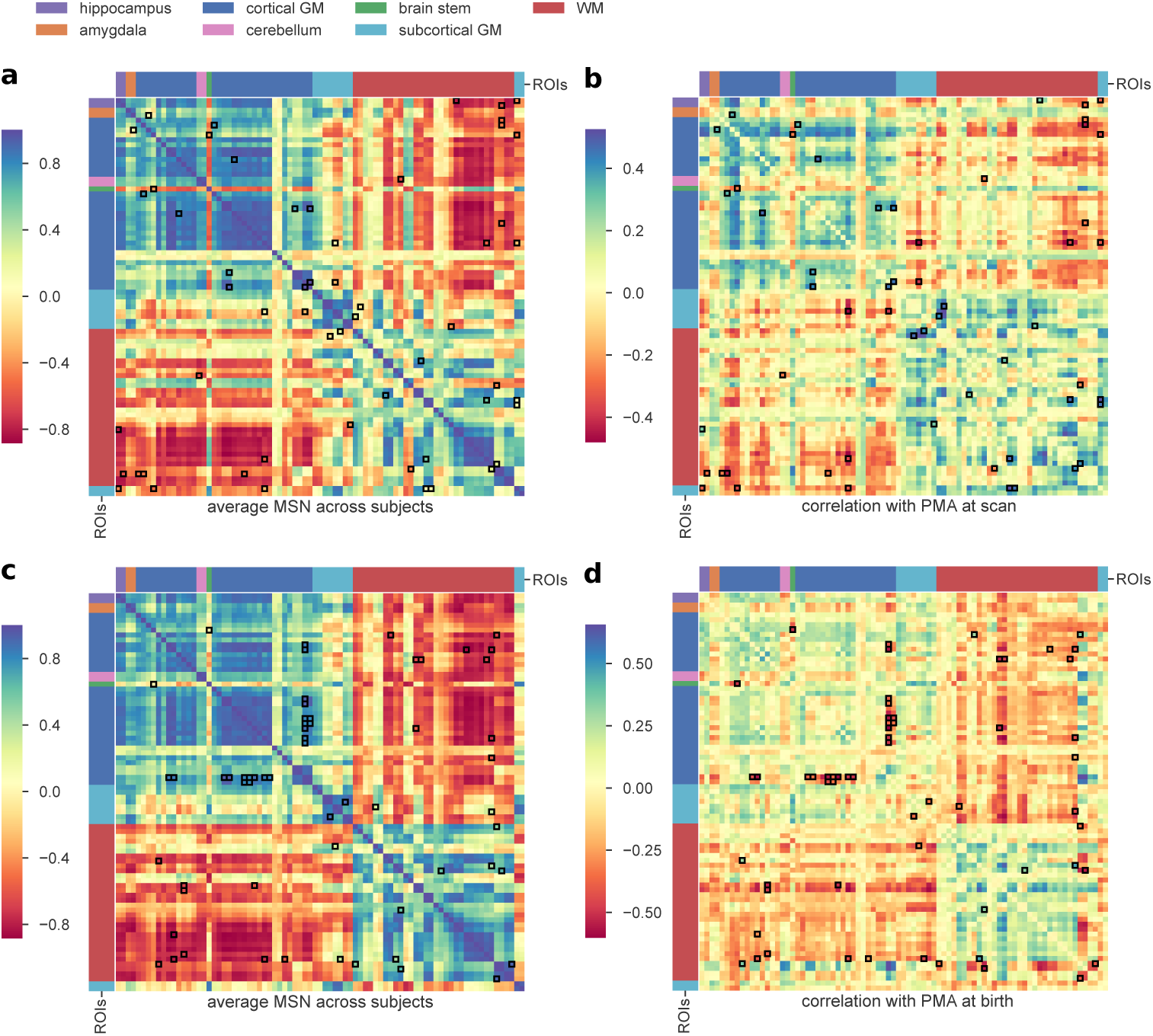
a) Average MSN computed across all subjects using the combination of features selected through the backward feature selection scheme for the age prediction task (Volume, FA, MD, AD, MK, *υ*_iso_, ODI_P_). b) Correlation between each connection weight (inter-regional similarity) shown in (a) and PMA at scan across subjects. c) Average MSN computed across all subjects using the combination of features selected through the backward feature selection scheme for the classification task (Volume, T1/T2, FA, MD, AD, RD, MK, *υ*_ic_, *υ*_iso_, ODI_P_, ODI_TOT_). d) Correlation between each connection weight (inter-regional similarity) shown in (c) and PMA at birth across subjects. Connections that were identified as predictive features by the models are highlighted in black. ROIs are ordered as in Supplementary Table S1.

The best classifier model was based on eleven out of the twelve features (all except ODI_S_), so compared to the age prediction model, four additional features were included: T1/T2, RD, *υ*_ic_ and ODI_TOT_. The average MSN computed with the selected features and the matrix of correlation with PMA at birth is shown in Fig. 5 (panels b and c). Comparing panel b and d of Fig. 5, it is apparent that while the patterns of correlation with PMA at scan and at birth are similar within GM and WM, subcortical ROIs show an opposite trend: with increasing PMA at scan subcortical ROIs tend to become more similar to WM ROIs and more dissimilar to GM ROIs, but the similarity between subcortical ROIs and cortical GM is positively correlated to age at birth.

### 3.2. Prediction results

The best regression model selected with LOOCV predicted chronological age (PMA at scan) with a MAE of 0.70 *±* 0.56 weeks on the test data, and a correlation between the predicted and the actual age equal to *r* = 0.78 (*p* = 1.71 *×* 10*^−^*^22^) (Supplementary Fig. S5). The results of the permutation test are shown in Fig. 6 and Supplementary Fig. S6. The confounding variables (gender and age at birth) were not selected by the internal feature selection procedure, hence the predictions were based on network features alone. To test whether there was any systematic difference in the predicted age between the term and the preterm group, we compared the error distributions with a Wilcoxon rank-sum test, but the result was not significant (*W* = 1108*, p* = 0.1085). For comparison, we evaluated the predictive performance of a linear regression model using only gender and PMA at birth as independent variables, that achieved a MAE of 1.03 *±* 0.88 weeks. A Wilcoxon signed-rank test confirmed that the latter model achieved a significantly greater error (*W* = 1633*, p* = 0.0001). Also models based on single global metrics and single-modality MSNs models provided poorer predictive performance than the selected multi-modality MSNs model (brain volume: MAE= 0.93 *±* 0.68, *R* = 0.58; median FA: MAE= 0.88 *±* 0.63, *R* = 0.58; structural: MAE= 1.08 *±* 0.79, *R* = 0.32; DKI: MAE= 0.94 *±* 0.70, *R* = 0.57; NODDI: MAE= 0.88 *±* 0.69, *R* = 0.61) and this was confirmed by a Wilcoxon signed-rank test (brain volume: *W* = 1813*, p* = 0.0019; median FA: *W* = 2045*, p* = 0.0184; structural: *W* = 1361*, p* = 2.76 *×* 10*^−^*^06^; DKI: *W* = 1734*, p* = 0.0004; NODDI: *W* = 1811*, p* = 0.0009). Conversely, the baseline model based on the ensemble on individual metrics used to build the best performing MSN model achieved similar performances (MAE: 0.72*±*0.56, *R* = 0.77). A scatter plot of the residuals of the two models (Supplementary Fig. S11) showed a linear trend, indicating that the two models share a similar information content.

**Figure 6:**
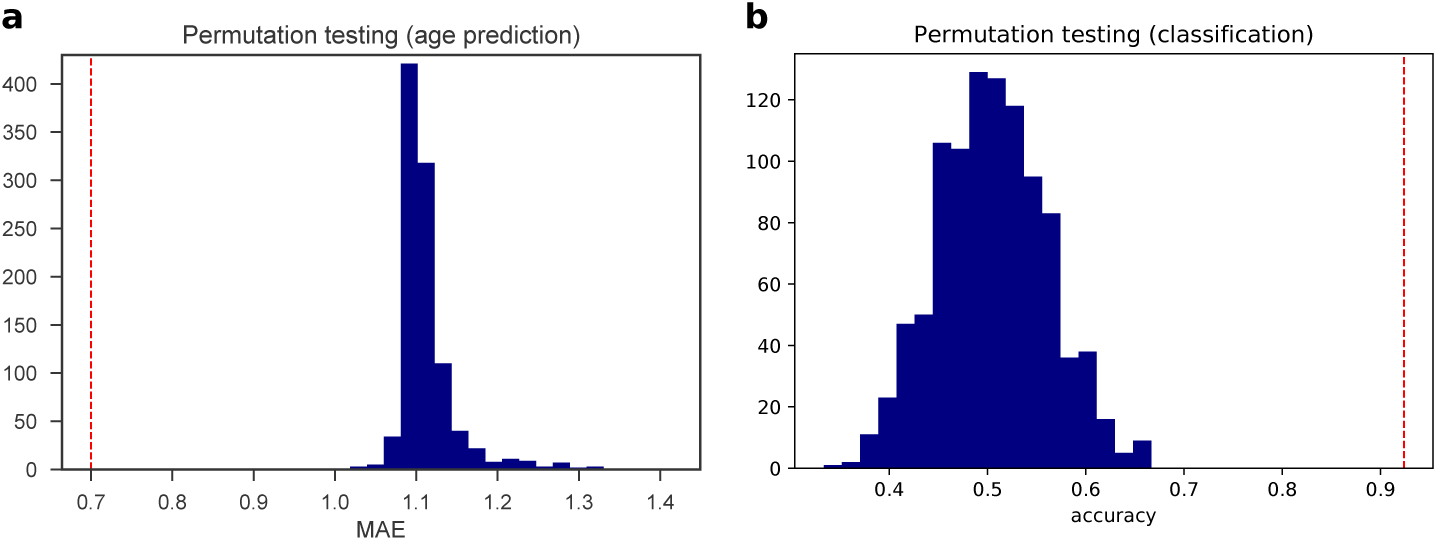
Null distributions computed over 1000 random permutations of the target variable for the age prediction (a) and the classification tasks (b). The red dotted lines indicate the performances of our models.

Supplementary Fig. S2 shows the results computed with 10-5-fold cross-validation in. All compared models performed similarly under the 10-5-fold scheme, and in general worse than with the LOOCV scheme, with the selected model achieving a MAE of 1 *±* 0.2 weeks (Supplementary Fig. S7).

To study which connections contributed the most to chronological age prediction, we selected only edges which were assigned a non-zero coefficient in at least 99% of cross-validation folds. These edges are shown in the chord diagram in Fig. 7 (realised with Circos, Krzywinski et al. (2009)), and are colour coded to distinguish between inter-regional similarities that increase or decrease with age, to highlight networks of regions whose morphological properties are converging (gray) or that tend to differentiate with increasing age (red). Intuitively, these edges connect ROIs whose anatomical and micro-structural properties are changing more than others between 38 and 45 weeks PMA, making the ROIs more or less similar. In other words, it is the relative timing of maturation of different brain tissues to determine the relevance of a connection in the age prediction task. The selected connections are located in both cortical (frontal, temporal, parietal and occipital lobes; insular and posterior cingulate cortex) and subcortical regions (thalamus, subthalamic and lentiform nuclei), in the brain stem and in the cerebellum. These areas have been previously associated with age-related changes and preterm birth (Boardman et al., 2006; Ball et al., 2013a; Batalle et al., 2017). For comparison, we report in Supplementary Table S2 the regional metrics selected as most predictive of age in the baseline model based on individual metrics.

**Figure 7:**
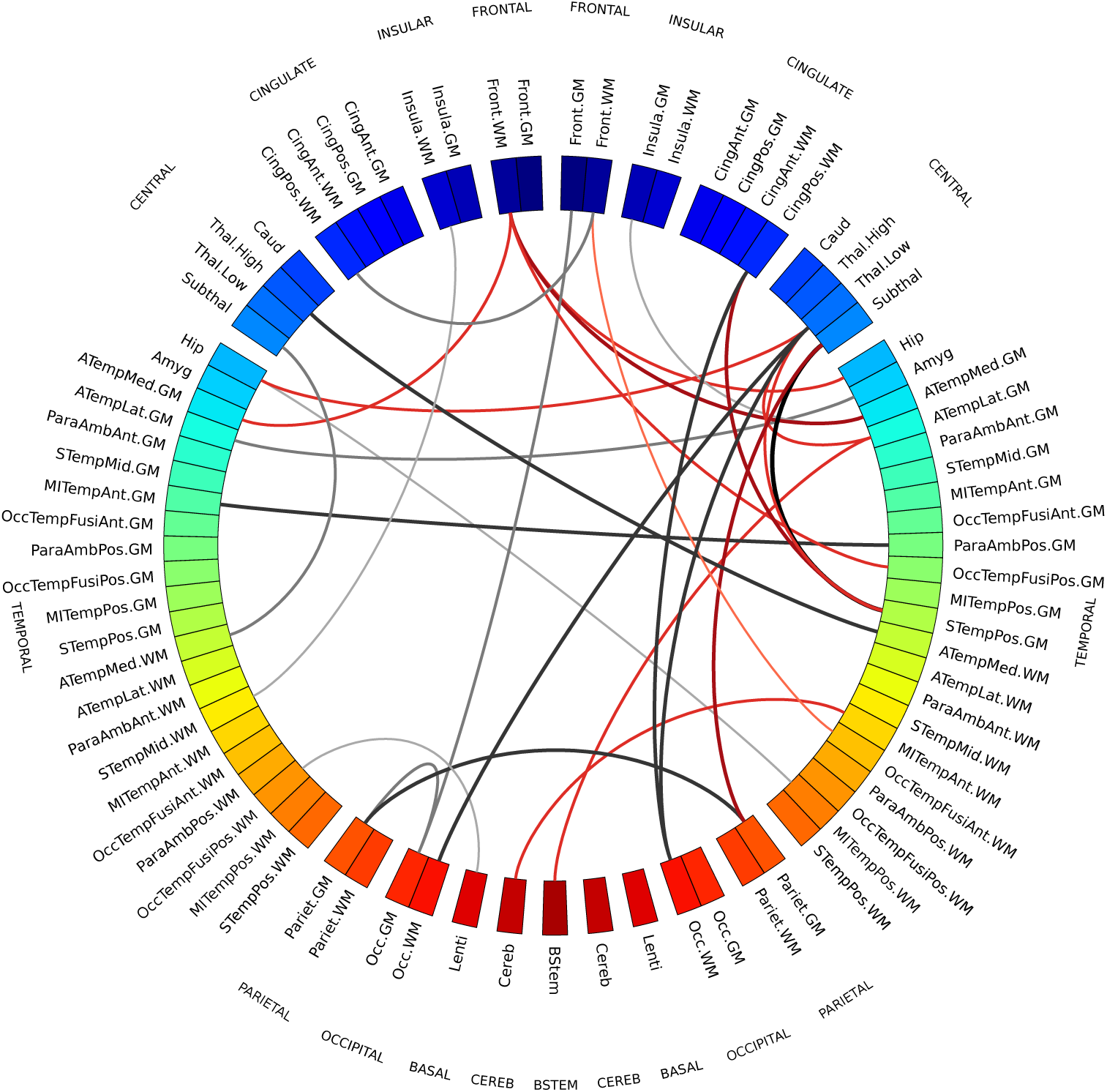
Chord diagram showing MSN edges used for age prediction in at least 99% of regression models in the cross-validation folds. Connections shown in gray are inter-regional similarities that increase with chronological age, while connections in red are inter-regional similarities that decrease with chronological age. The edge width is proportional to the correlation between inter-regional similarities and PMA. The left side of the diagram corresponds to the left side of the brain. Abbreviations for ROI names are explained in Supplementary Table S1.

**Table 2:**
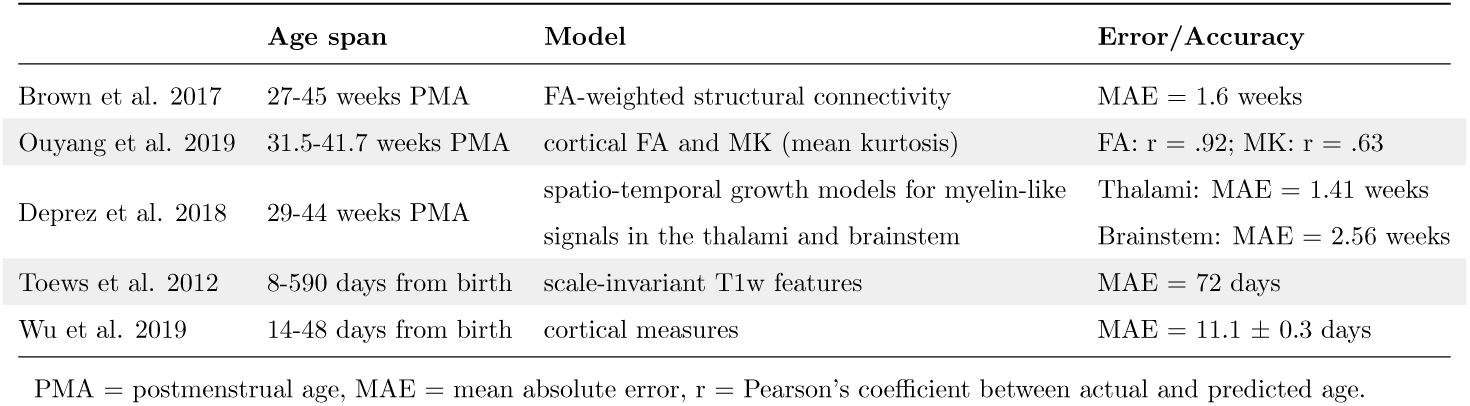
Results from previous works in the age prediction task.

The best classifier discriminated between term and preterm infants with a 92% LOOCV accuracy (Fig. 6). None of the confounders were included among the selected features. A logistic regression model built on age at scan and gender did not achieve significant accuracy (56%*, p* = 0.091), while adding motion to the predictors produced a 61% accuracy, slightly above chance level (*p* = 0.03), but it should be noted that a model based on motion only was 59% accurate (*p* = 0.02). Models based on global features achieved 55% accuracy for total brain volume and 56% accuracy for median FA. Models built on single data modalities attained 65% accuracy for structural features only, 89% accuracy for DKI features only, and 88% accuracy for NODDI features only. Results computed with 10-5-fold cross-validation are shown in Supplementary Fig. S4. The best classifier selected with LOOCV also achieved top accuracy with 10-5-fold (accuracy 90%, Supplementary Fig. S7).

The network of regions that showed the most divergent pattern of structural brain properties in preterm versus term infants comprised the brain stem, the thalamus and the subthalamic nucleus; WM regions in the frontal and insular lobes; GM regions in the occipital lobe; both WM and GM regions in the temporal and parietal lobes and in the posterior cingulate cortex. The chord diagram of edges selected by 99% of the models is shown in Fig. 8, in red where inter-regional similarities are greater in the term group and in gray where they are greater in the preterm group. For comparison, Supplementary Table S3 lists the regional metrics selected by the baseline model based on individual metrics, that obtained a 94% accuracy.

**Figure 8:**
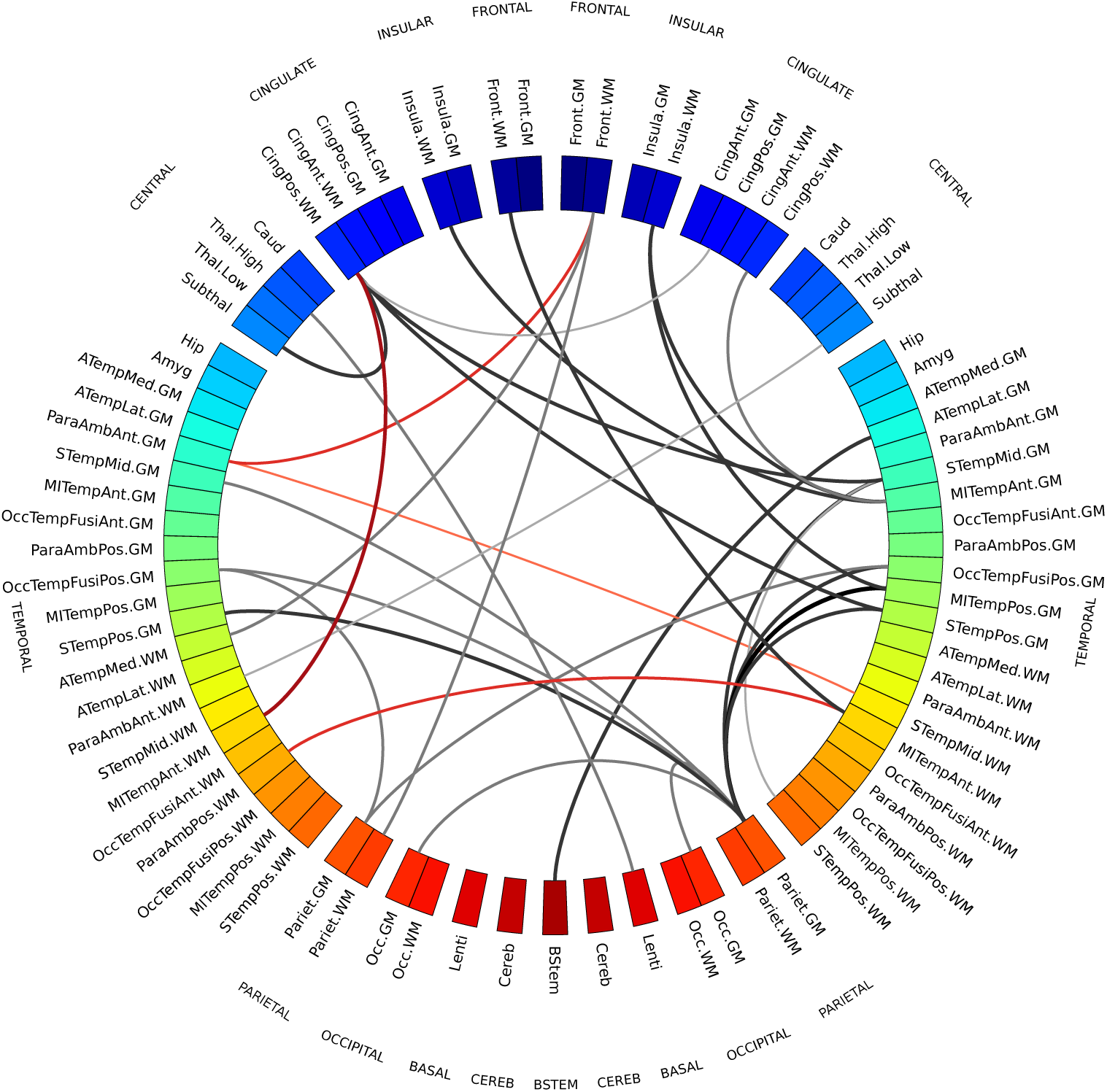
MSN edges showing a divergent pattern of morphological properties in term and preterm infants in at least 99% of classification models in the cross-validation folds. Gray connections indicate inter-regional similarities that are greater in the preterm group, while red connections are greater in the term group. The edge width is proportional to the correlation between inter-regional similarities and prematurity. The left side of the diagram corresponds to the left side of the brain. Abbreviations for ROI names are explained in in Supplementary Table S1.

### 3.3. Testing for asymmetry

In both chord diagrams (Figs. 7 and 8) we observed more edges in the right hemisphere than in the left one. Both elastic net and SVM models perform a feature selection step to exclude features that are correlated and that carry redundant information in order to improve prediction performance, hence it might be the case that the models selected the right connections and discarded the left ones precisely because they had a similar information content. Additionally, in the leave-one-out cross-validation scheme the training sets only differ by two samples in each fold, hence models might be similar across folds.

To test the hypothesis that the two hemispheres carry a different information content, we performed two experiments. First, we repeated the same analyses extracting inter-regional similarities from either the right or the left hemisphere. We compared the performance obtained with the regression and classification models on the different subsets of features used in the backward feature selection scheme in the main analyses. We found that for the age prediction model a Wilcoxon signed-rank test testing the hypothesis that the prediction error was higher using only connections from the left hemisphere was significant (*W* = 156*, p* = 2.57 *×* 10*^−^*^11^), while there was no statistically significant difference in the case of the classification model. These results replicated also when using 10-5-fold cross-validation (age prediction: *W* = 160*, p* = 2.98 *×* 10*^−^*^11^; no significant difference in classification). We also compared the residuals obtained using either the right or the left hemisphere for age prediction with the set of features selected with backward feature selection (Supplementary Fig. S11) and found that the residuals of the fitted models are linearly correlated, suggesting that the two hemispheres do carry a similar information content, but one presents clearer signal than the other. We then used permutation testing to test the “interchangeability” of right and left regions: starting from the subsets of imaging metrics selected in the main analyses for the age prediction and classification models, we generated two null distributions by randomly swapping a subset of homotopic brain regions between the right and left hemisphere, and then repeating the exact same analyses 1000 times. We then counted how many times in the random models there was a disproportion of inter-regional similarities selected in the right hemisphere equal or greater than the one we observed with our models. If the right and left are “interchangeable”, the number of interregional similarities selected should remain the same on average. We found that in the age prediction task, under the null distribution, the disproportion of predictive connections in the right hemisphere was associated with a *p* = 0.036, while in the classification task the disproportion was not significant (*p* = 0.166). This implies that at least for age prediction the two hemispheres are not inter-changeable, suggesting again that the right hemispheres has a stronger signal. A similar trend was observed under the 10-5-fold cross-validation scheme, but in this case we could not reject the null hypothesis that inter-regional similarities are selected with the same frequency from both hemispheres (*p* = 0.098).

## 4. Discussion

These results show that the information encoded in MSNs is predictive of chronological brain age in the neonatal period and that MSNs provide a novel data-driven method for investigating neuroanatomic variation associated with preterm birth. MSNs were built by combining features from different imaging sequences that describe complementary aspects of brain structure that have been previously studied in isolation (Makropoulos et al., 2016; Batalle et al., 2017) and the resulting predictive models achieved a high accuracy for age prediction and classification. By comparing the performance of MSNs features with basic demographic information (age at birth and gender) and simple metrics such as total brain volume and median white matter FA, we also showed that integrating imaging data provides relevant additional information to characterise brain age. Although we cannot exclude the possibility that some of the variability shared with age at birth, gender or brain volume is encoded in the imaging variables, the comparative analysis and the permutation testing results showed that the observed variance cannot be completely explained by demographic variables or simpler metrics alone. However, a high accuracy is not the only goal of the proposed method: once we have determined that the model is able to learn a relationship between the MSN features and age or prematurity, we can interrogate it to find out which features, regions and structures are involved in the predictions, thus allowing for further inferences.

We anticipate that the main clinical and research utilities of MSNs will be to investigate divergent maturational patterns in the context of perinatal environmental, genetic and clinical exposures, leading to improved understanding of antecedents to, and consequences of, atypical brain development. For these purposes a prediction tool with average 5 days error is highly precise compared with other methods for assessing brain maturation, which usually rely upon simple linear regression, use single image features, or broad classifications of prematurity (Toews et al., 2012; Brown et al., 2017; Batalle et al., 2018; Deprez et al., 2018; Bouyssi-Kobar et al., 2018; Ouyang et al., 2018).

The regions identified as most predictive have been previously associated with age-related changes and preterm birth (Boardman et al., 2006; Ball et al., 2013a; Batalle et al., 2017; Bouyssi-Kobar et al., 2018). These data suggest that to fully describe morphological variation in the developing brain it may be advantageous to adopt a holistic approach, leveraging the additional information that can be derived from integrating multi-contrast MRI data. The main motivation for using a network-based approach is to obtain a whole-brain description of a developmental pattern. By using topologically integrated features instead of single metrics it is possible to access an additional layer of information that is not explicitly encoded in the individual metrics, i.e. how the relationships between metrics vary in different parts of the brain. Working with correlations instead of an ensemble of heterogeneous metrics also aids interpretation, as the focus is shifted from the values of single metrics across the brain, each influenced by disparate factors, to similarities between brain regions, which is a more relatable concept. Additionally, the adoption of a network model has proven to be a useful abstraction to capture the modular organisation of the brain: in the original work introducing MSNs to study microscale cortical organization in adults, the authors demonstrated that regions that were similar in MSNs were more likely to belong to the same cytoarchitectonic class, to be axonally connected and to have high levels of co-expressions of genes specialised for neural functions (Seidlitz et al., 2018). Another reason for working with similarities instead of single regional metrics is methodological: computing edge weights as inter-regional similarities enables an integrated representation of several metrics in a single network; to work with the original features directly would mean either working with several networks (thus requiring a further step to integrate them and aggravating the problems related with the “curse of dimensionality”) or concatenating all the features in a single predictive model (thus excluding the interactions between metrics from the model).

Our data are consistent with previous studies of perinatal brain age prediction based on a single type of data or a single metric. For example, Brown et al. (2017) used dMRI tractography to predict brain dysmaturation in preterm infants with brain injury and abnormal developmental outcome and found that altered connectivity in the posterior cingulate gyrus and the inferior orbitofrontal cortex were associated with a delayed maturation; both of these regions are included in the networks identified by our model. Regional FA, MD, MK, and *υ*_ic_ are each predictive of age (Genc et al., 2017; Karmacharya et al., 2018; Ouyang et al., 2019), and the first three measures were selected in our age predicition model. Growth of the thalami and brainstem, defined in terms of myelin-like signals from T2-weighted images, successfully predicted age between 29 and 44 weeks (Deprez et al., 2018) and these regions are included in the networks most predictive of age in the current study. In Toews et al. (2012), scale-invariant image features were extracted from T1-weighted MRI data of 92 subjects over an age range of 8-590 days to build a developmental model that was used to predict age of new subjects; and Ceschin et al. (2018) proposed a deep learning approach to detect subcortical brain dysmaturation from T2-weighted fast spin echo images in infants with congenital hearth disease. Wu et al. (2019) used cortical features extracted from structural images to predict age of 50 healthy subjects with 251 longitudinal MRI scans from 14 to 797 days; in accordance with our results, the regions reported to be important for age prediction were bilateral medial orbitofrontal, parahippocampal, temporal pole, right superior parietal and posterior cingulate cortex. Although our results are not directly comparable with the above works because of the heterogeneity of employed models, validation techniques and population variation (different age ranges), our prediction error is among the lowest reported (see Table 2 for a summary of previous results), but it should be noted that there is a strong positive correlation between the reported MAEs and the age range of the samples. In addition, many works have identified imaging biomarkers associated with preterm birth, such as brain tissue volume (Alexander et al., 2018; Gui et al., 2019), myelin content (Melbourne et al., 2016), and diffusion tensor metrics (Anjari et al., 2007; Bouyssi-Kobar et al., 2018).

The connections most predictive of age revealed that brain maturation is characterised by morphological convergence of some networks and divergence of others (Fig. 7). These connections mostly involve fronto-temporal and subcortical ROIs, which suggests that the micro- and macro-structural properties of these regions are highly dynamic between 38-45 weeks. Among these, interregional similarities within GM and WM increase with age, similarities between cortical GM and WM decrease, while subcortical ROIs become more similar to WM and more dissimilar to cortical GM. This is consistent with previous findings on the different trends in development of the thalamus and the cortex (Eaton-Rosen et al., 2015). Additionally, in a study of early development of structural networks (Batalle et al., 2017), connections to and from deep grey matter are reported to show the most rapid developmental changes between 25-45 weeks, while intra-frontal, frontal to cingulate, frontal to caudate and inter-hemispheric connections are reported to mature more slowly.

Conversely, the inter-regional similarities selected by the SVM classifier to discriminate between term and preterm (Fig. 5) are more distributed across cortical GM and WM and are for the most part greater in the preterm group. The fact that in the term group these cortical ROIs are less homogeneous in terms of structural properties could be interpreted as a sign that in term infants these regions are at a different stage of maturation where their morphological profile is consolidating along specialised developmental trajectories. It has been previously suggested that the rapid maturation of cortical structures occurring in the perinatal period is vulnerable to the effects of preterm birth (Kostović and Jovanov-Milošević, 2006; Ball et al., 2011, 2013b; Smyser et al., 2016b).

The differences between networks identified for age prediction and for preterm classification indicate that atypical brain development after preterm birth is not solely a problem of delayed maturation, but it is characterised by a specific signature. Indeed, while the age prediction networks capture changes occurring in both the preterm and the term group, the classification networks highlight where there are group-wise differences, and they do not match: in the case of a delayed maturation we would have observed differences in the same regions undergoing age-related changes. MSN variations associated with preterm birth affected brain stem, thalami, sub-thalamic nuclei, WM regions in the frontal and insular lobes, GM regions in the occipital lobe, and WM and GM regions in the temporal and parietal lobes and in the posterior cingulate cortex. This distribution of structural variation is consistent with previous reports of regional alteration in brain volume and dMRI parameters based on single contrasts (Boardman et al., 2006; Bonifacio et al., 2010; Ball et al., 2013a; Brown et al., 2017; Batalle et al., 2017; Alexander et al., 2018; Thompson et al., 2018b; Bouyssi-Kobar et al., 2018). Furthermore, compared to the age prediction model, the MSNs used for preterm classification are based on four additional metrics: T1/T2, related to myelination; RD, measuring water dispersion; *υ*_ic_ describing neurite density; and ODI_TOT_, associated with the fanning of WM tracts. All these metrics contribute to characterise the micro-structural alterations associated with preterm birth (Eaton-Rosen et al., 2015; Melbourne et al., 2016; Batalle et al., 2018; Thompson et al., 2018b; Bouyssi-Kobar et al., 2018).

We observed a disproportion in the distribution of the connections selected by our models, with a preference for the right hemisphere, hinting at the existence of lateralization in the maturational process. An asymmetry in the development of the right hemisphere in neonates was previously reported in Dubois et al. (2010); Yap et al. (2011); Wu et al. (2019), and our experiments (section 3.3) partially supported the hypothesis that the right hemisphere plays a relevant role in the context of age prediction.

### 4.1. Limitations

This work has some limitations. First, compared with the original work on MSNs (Seidlitz et al., 2018), we did not have a multi-parametric mapping sequence (Weiskopf et al., 2013); however, because the model is extensible, information from other contrasts could be added and evaluated for their effect on prediction. The MSN model could also be applied to study the properties of cortical gray matter (such as thickness, sulcal depth or curvature), that have been previously reported to be predictive of age in children (Brown et al., 2012) and could contribute significantly in characterising the newborn brain. However, metrics that only apply to selected structures (e.g. the cortex) cannot be used in a whole brain analysis, as to compute inter-regional similarities each region needs to be described by the same set of metrics. This particular study was designed based on prior knowledge that typical development and atypical development associated with preterm birth are characterised by global changes (Ball et al., 2013a; Anderson, 2014; Eaton-Rosen et al., 2015; Melbourne et al., 2014), and MSNs integrating dMRI and sMRI data were chosen to study generalised processes across the whole brain.

Second, we used a motion correction technique that attenuates the impact of head motion on structural connectivity (Andersson and Sotiropoulos, 2016; Baum et al., 2018), and we found that scanner motion was not contributing significantly to prediction accuracy; however we cannot rule out a possible confounding effect of motion on the estimation of regional metrics.

Third, the preterm study population was representative of survivors of modern neonatal intensive care in terms of gestational age range and prevalence of co-morbidities of preterm birth that may influence brain maturation, but it is still possible that the results were influenced by biological variability specific to the cohort. A replication study will be required to determine whether the patterns of dysmaturation we found are generalisable.

Finally, we assessed the performance of our models with both LOOCV and 10-5-fold schemes in order to investigate the stability of our findings with respect to the chosen cross-validation scheme and we observed some variability in the general trends of the results. The disagreement we found might derive from the limited size of the training set in the case of the repeated-5-fold scheme (all models tended to perform worse, suggesting there were not enough samples for learning), and this was indeed the reason why our first choice was the leaveone-out scheme. As it is always the case when working with machine learning, increasing the sample size would increase the power of the models, thereby reducing the margin of error and the risk of overfitting, with the result that both schemes should converge to similar findings.

### 4.2. Conclusions

Combining multiple imaging features in a single model enabled a detailed description of the morphological properties of the developing brain that was used inside a predictive framework to identify two networks of regions: the first, predominantly located in subcortical and fronto-temporal areas, that contributed most to age prediction: the second, comprising mostly frontal, parietal, temporal and insular regions, that discriminated between preterm and term born infant brains. Both predictive models performed best when structural, diffusion tensor-derived and NODDI metrics were combined, which demonstrates the importance of integrating different biomarkers to generate a global picture of the developing human brain. The achieved accuracy supports the hypothesis that studying the interaction between regional metrics can shed light on the mechanics of development.

Morphology, structural connectivity and maturation are all influenced by genetics, co-morbidities of preterm birth, and nutrition (Boardman et al., 2014; Anblagan et al., 2016; Sparrow et al., 2016; Krishnan et al., 2016; Ball et al., 2017; Alexander et al., 2018; Blesa et al., 2019). In future work MSNs could offer new understanding of the impact of these factors on integrated measures of brain development, and the relationship between neonatal MSNs and functional outcome could bring novel insights into the neural bases of cognition and behaviour, by identifying networks of regions associated with later development. MSNs could also enable a direct comparison with functional networks extracted from fMRI, to explore how structure and function interplay in the neonatal period, and study how well the two network models together explain individual variability in developmental outcome.

## Acknowledgements

We are grateful to the families who consented to take part in the study. This work was supported by Theirworld (www.theirworld.org) and was undertaken in the MRC Centre for Reproductive Health, which is funded by MRC Centre Grant (MRC G1002033). MJT was supported by NHS Lothian Research and Development Office. Participants were scanned in the University of Edinburgh Imaging Research MRI Facility at the Royal Infirmary of Edinburgh which was established with funding from The Wellcome Trust, Dunhill Medical Trust, Edinburgh and Lothians Research Foundation, Theirworld, The Muir Maxwell Trust and many other sources; we thank the University’s imaging research staff for providing the infant scanning.

