## Supplementary material for "Neonatal morphometric similarity mapping for predicting brain age and characterizing neuroanatomic variation associated with preterm birth"

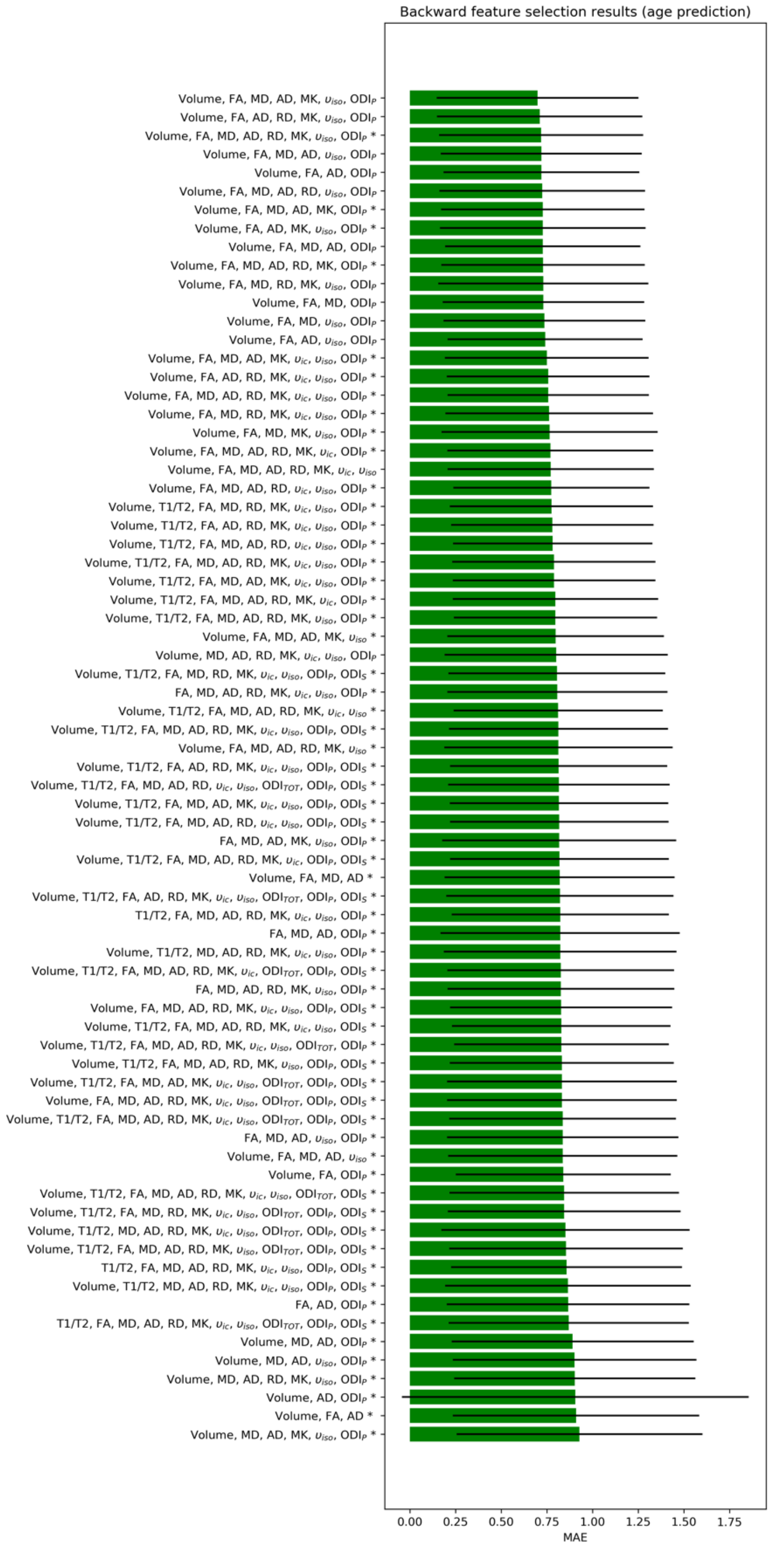


*Figure S1 Feature selection results for the age prediction task. The best set of features was selected with a backward feature selection scheme: starting from the full set of features, at each iteration the feature whose subtraction caused the least increase in prediction error was removed. The mean absolute error (MAE) computed with leave-one-out cross-validation is reported for each subset of features. The black lines depict standard deviation. Solutions marked with a * are significantly worse than the selected model (Wilcoxon signed-rank test, p<0.05)*

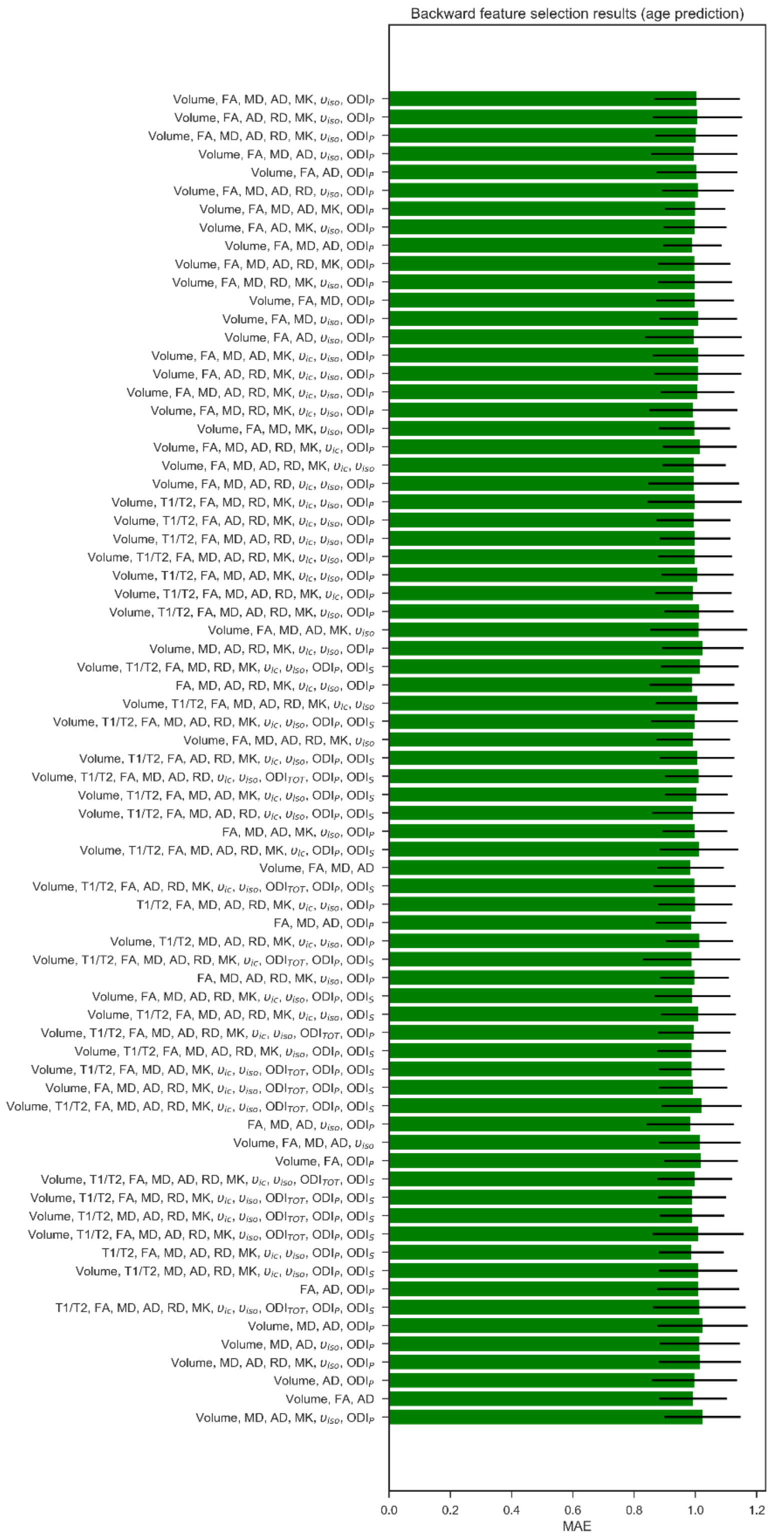


*Figure S2 The mean absolute error (MAE) computed with 10 repeated 5-fold cross-validation is reported for each subset of features in S1. The black lines depict standard deviation.*

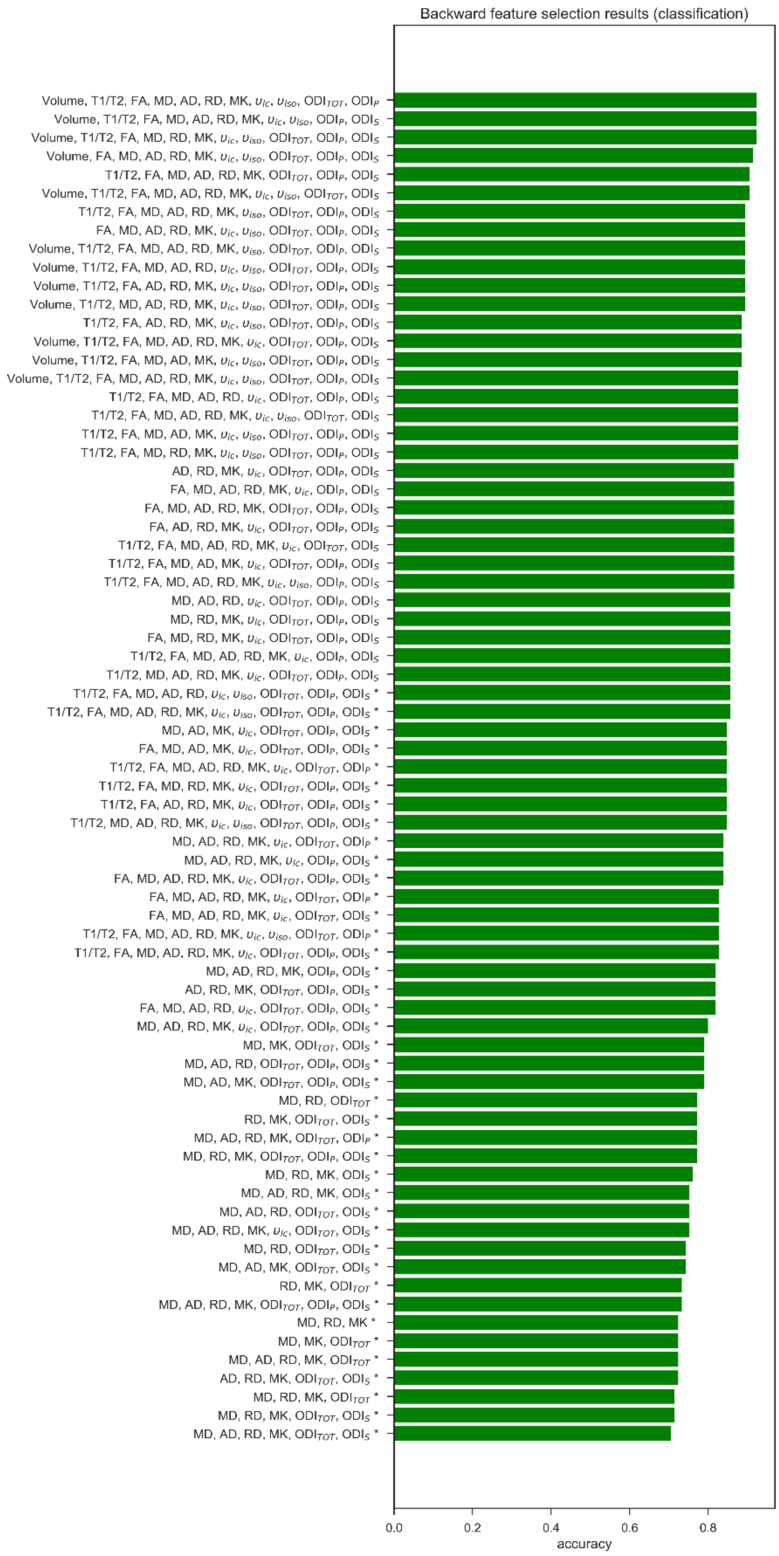


*Figure S3 Feature selection results for the classification task. The best set of features was selected with a backward feature selection scheme: starting from the full set of features, at each iteration the feature whose subtraction caused the least dencrease in prediction accuracy was removed. The accuracy computed with leave-one-out cross-validation is reported for each subset of features. Solutions marked with a * are significantly worse than the selected model (McNemar test, p<0.05)*

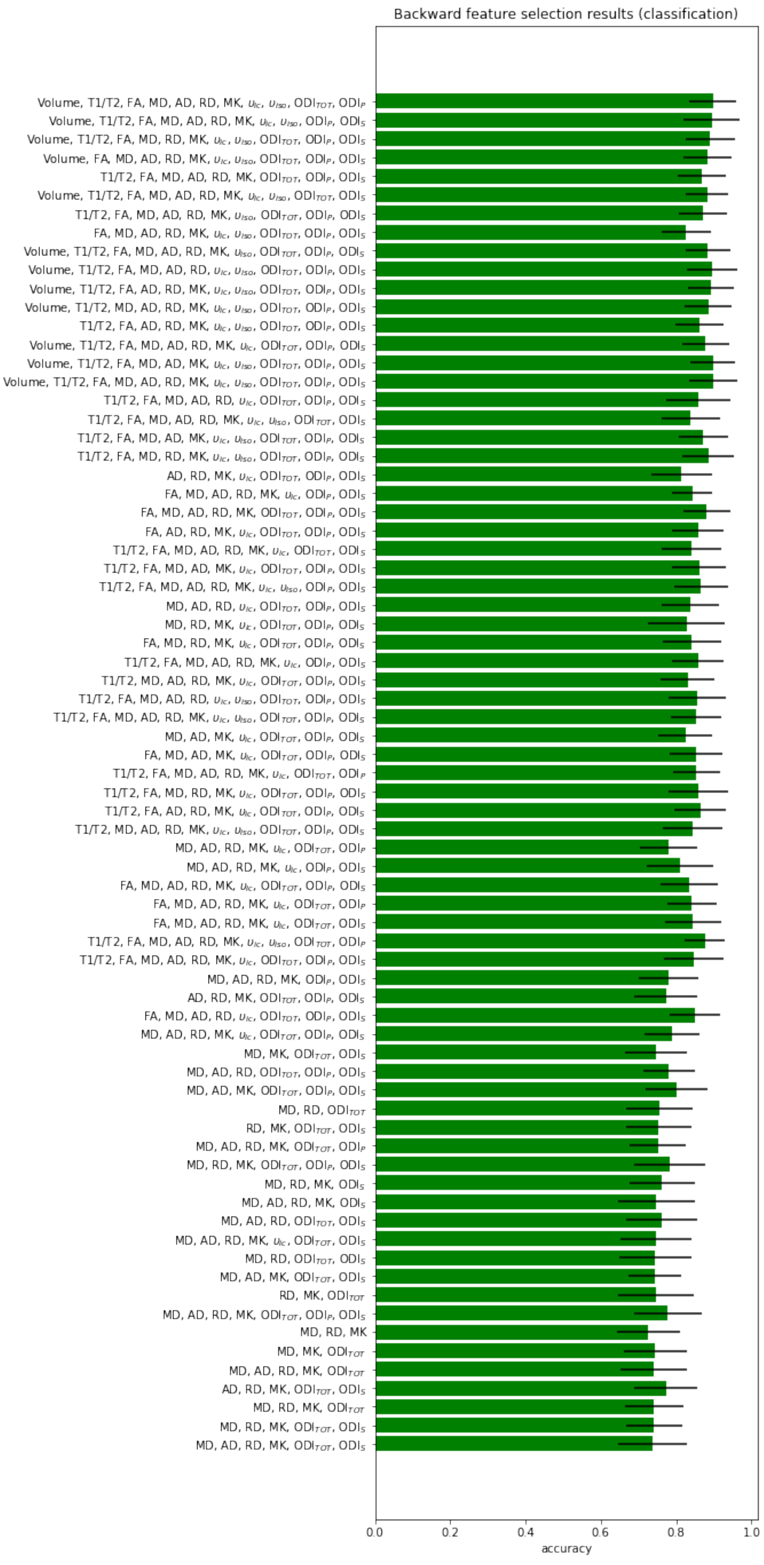


*Figure S4 The mean accuracy computed with 10 repeated 5-fold cross-validation is reported for each subset of features in S3. The black lines depict standard deviation across repetitions.*

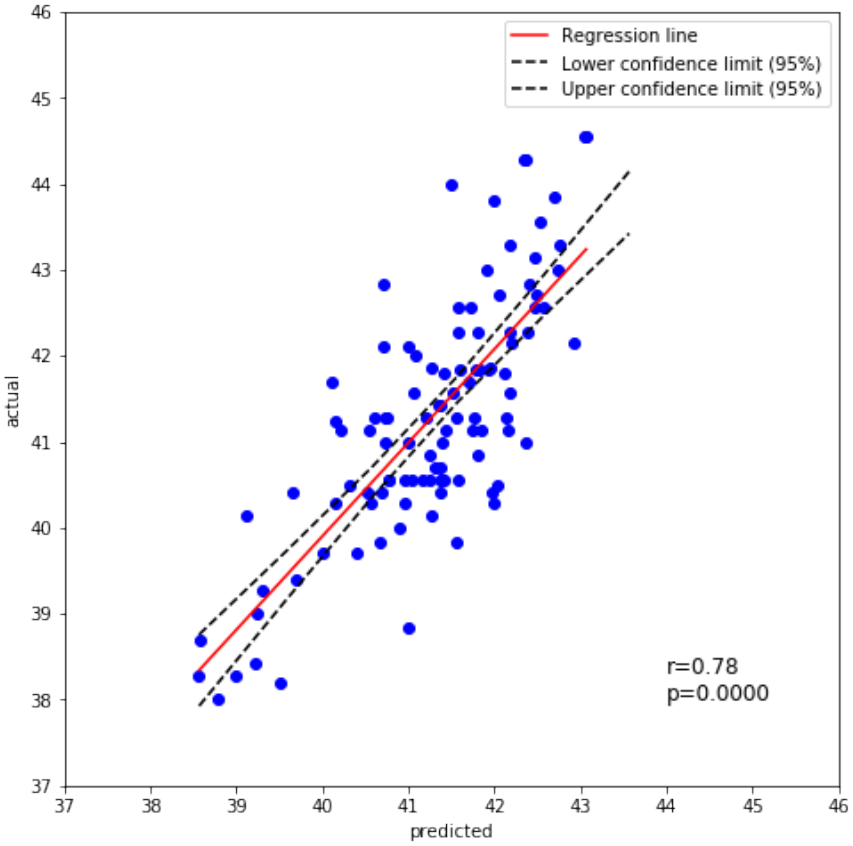


*Figure S5 Regression plot showing the correlation between actual and predicted age.*

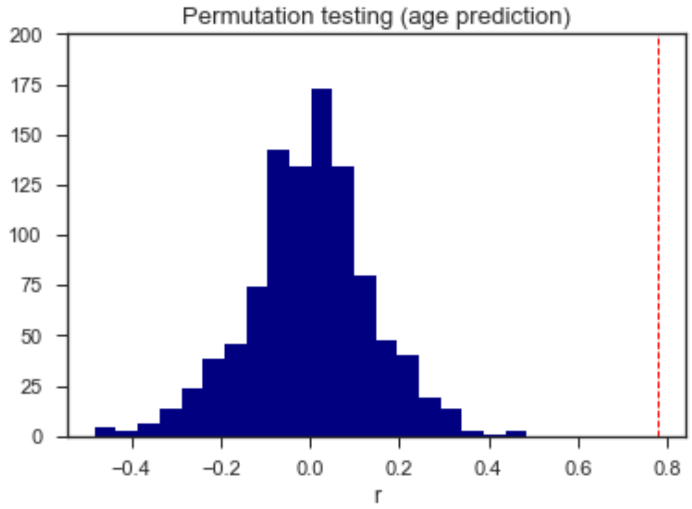


*Figure S6 Null distributions for the correlation between actual and predicted age (r) computed over 1000 random permutations of the target variable. The red dotted line indicates the performances of the selected model.*

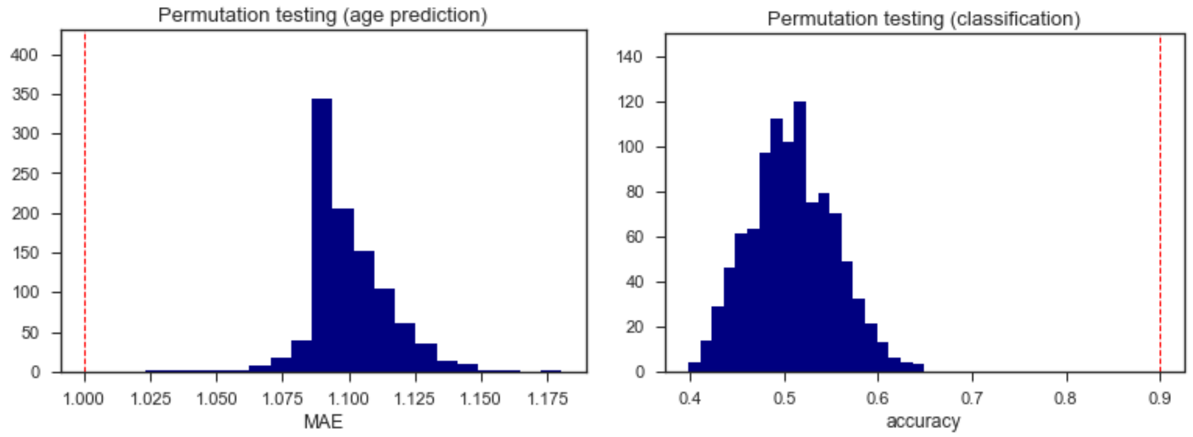


*Figure S7 Null distributions computed with 10-5-fold cross-validation over 1000 random permutations of the target variable for the age prediction task (left) and the classification task (right). The red dotted lines indicate the performances of the selected models.*

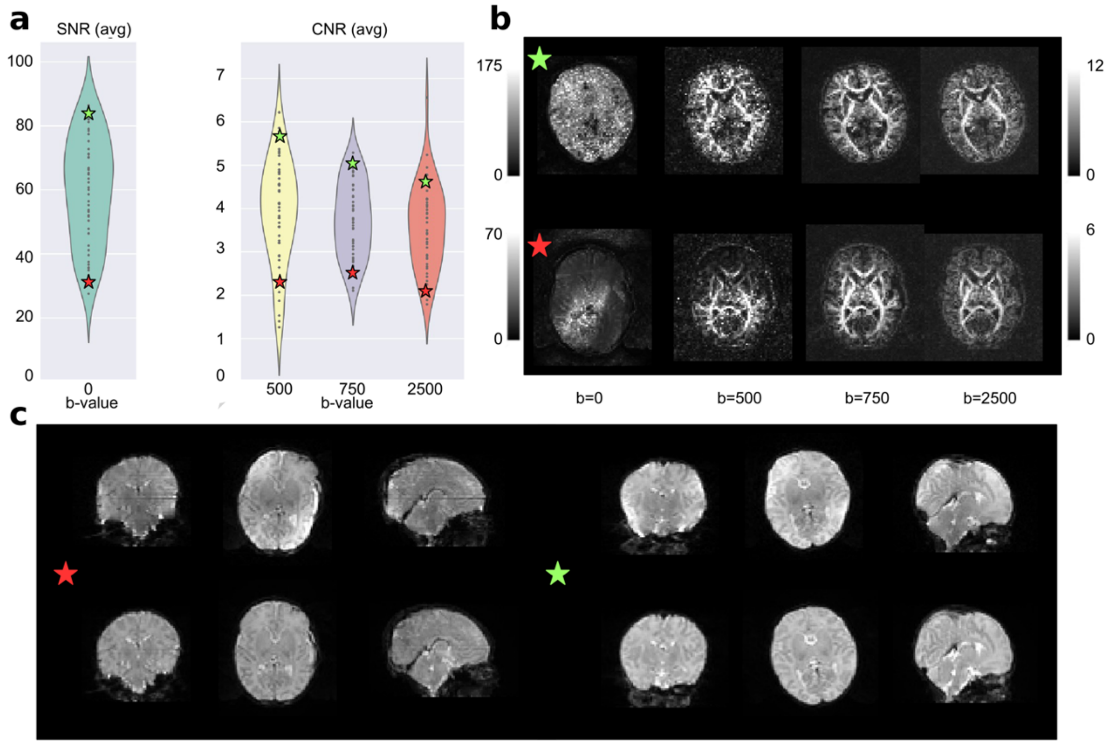


*Figure S8 Quality control results for the term population only. a) Results for the overall population with two selected subjects, one from the top quartile of the SNR and CNR distributions (green star) and the other from the bottom quartile (red star). b) The SNR and CNR maps for the selected subjects. c) The b0 of both subjects before and after the processing pipeline.*

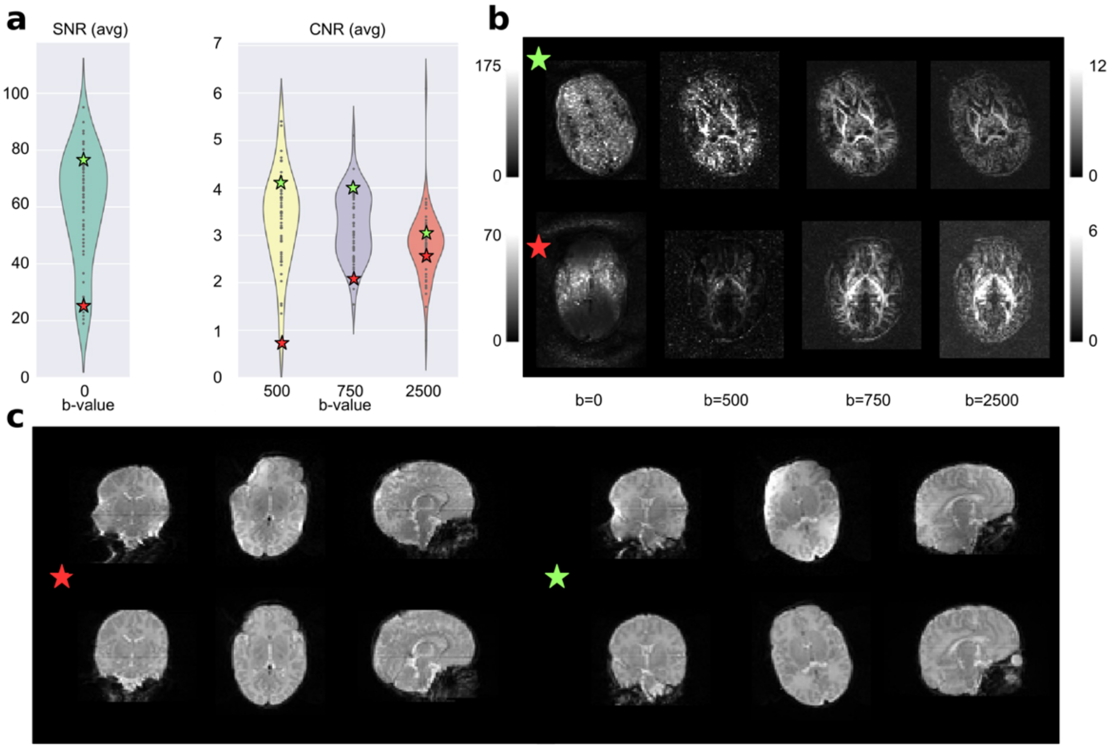


*Figure S9 Quality control results for the term population only. a) Results for the overall population with two selected subjects, one from the top quartile of the SNR and CNR distributions (green star) and the other from the bottom quartile (red star). b) The SNR and CNR maps for the selected subjects. c) The b0 of both subjects before and after the processing pipeline.*

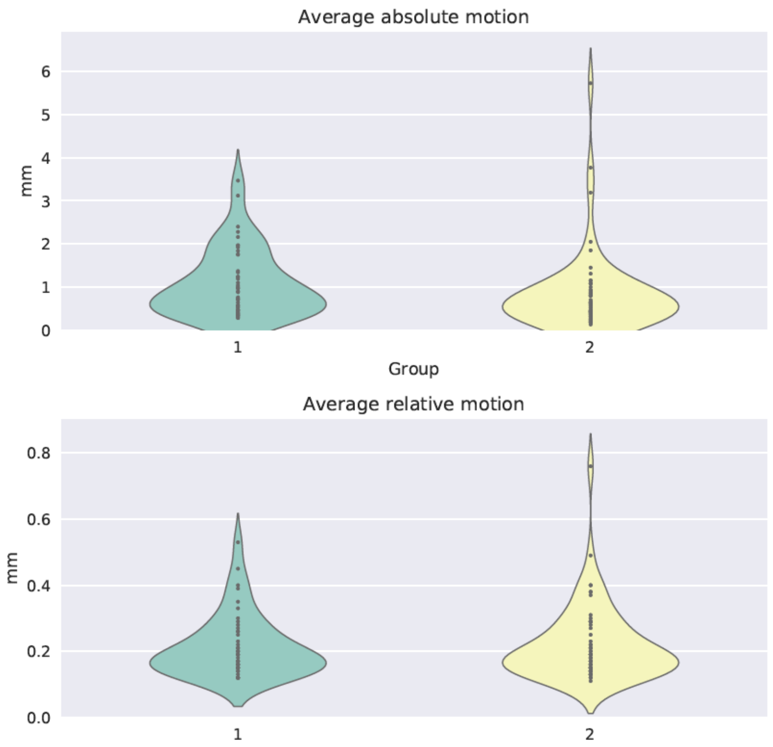


*Figure S10 Average absolute motion (computed as 3 translations and 3 rotations around the x, y and z axes, with respect to the reference volume) and average relative motion (same as before, but with respect to the previous volume) of the two different groups: term (1) and preterm (2).*

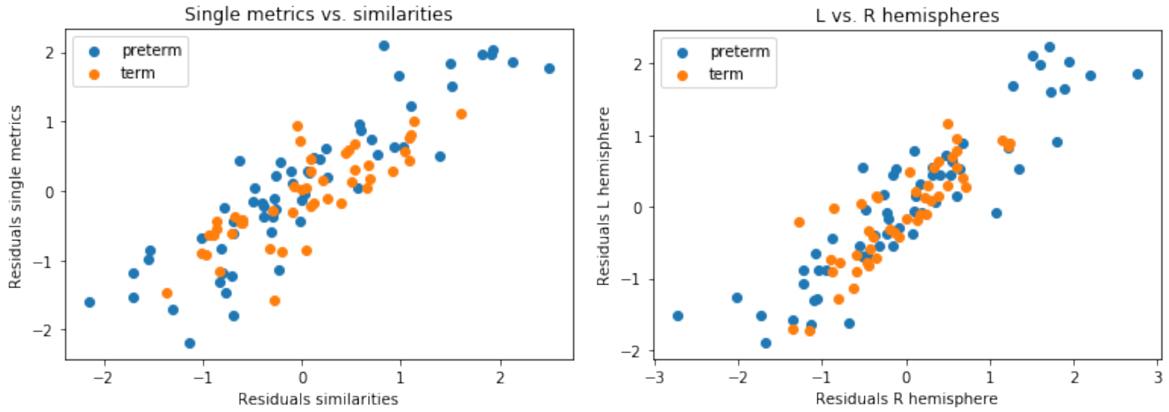


*Figure S11 (left) Scatter plot of the residuals computed with the baseline model (individual metrics) vs. the residuals computed with the MSN model for the set of features selected with backward feature selection. (right) Scatter plot of the residuals computed with a MSN model built on the left hemisphere only vs. the residuals computed with a MSN model built on the right hemisphere only, using the features selected with backward feature selection.*

*Table S1 Abbreviations of ROI names and division of ROIs into groups.*

| Short name | Long name | Group |
|---|---|---|
| Hip | Hippocampus left | hippocampus |
| Hip | Hippocampus right | hippocampus |
| Amyg | Amygdala left | amygdala |
| Amyg | Amygdala right | amygdala |
| ATempMed.GM | Anterior temporal lobe | cortical GM |
| ATempMed.GM | Anterior temporal lobe | cortical GM |
| ATempLat.GM | Anterior temporal lobe | cortical GM |
| ATempLat.GM | Anterior temporal lobe | cortical GM |
| ParaAmbAnt.GM | Gyri parahippocampalis et ambiens anterior part left GM | cortical GM |
| ParaAmbAnt.GM | Gyri parahippocampalis et ambiens anterior part right GM | cortical GM |
| STempMid.GM | Superior temporal gyrus | cortical GM |
| STempMid.GM | Superior temporal gyrus | cortical GM |
| MITempAnt.GM | Medial and inferior temporal gyri anterior part left GM | cortical GM |
| MITempAnt.GM | Medial and inferior temporal gyri anterior part right GM | cortical GM |
| OccTempFusiAnt.GM | Lateral occipitotemporal gyrus | cortical GM |
| OccTempFusiAnt.GM | Lateral occipitotemporal gyrus | cortical GM |
| Cereb | Cerebellum left | cerebellum |
| Cereb | Cerebellum right | cerebellum |
| BStem | Brainstem | brain stem |
| Insula.GM | Insula right GM | cortical GM |
| Insula.GM | Insula left GM | cortical GM |
| Occ.GM | Occipital lobe right GM | cortical GM |
| Occ.GM | Occipital lobe left GM | cortical GM |
| ParaAmbPos.GM | Gyri parahippocampalis et ambiens posterior part right GM | cortical GM |
| ParaAmbPos.GM | Gyri parahippocampalis et ambiens posterior part left GM | cortical GM |
| OccTempFusiPos.GM | Lateral occipitotemporal gyrus | cortical GM |
| OccTempFusiPos.GM | Lateral occipitotemporal gyrus | cortical GM |
| MITempPos.GM | Medial and inferior temporal gyri posterior part right GM | cortical GM |
| MITempPos.GM | Medial and inferior temporal gyri posterior part left GM | cortical GM |
| STempPos.GM | Superior temporal gyrus | cortical GM |
| STempPos.GM | Superior temporal gyrus | cortical GM |
| CingAnt.GM | Cingulate gyrus | cortical GM |
| CingAnt.GM | Cingulate gyrus | cortical GM |
| CingPos.GM | Cingulate gyrus | cortical GM |
| CingPos.GM | Cingulate gyrus | cortical GM |
| Front.GM | Frontal lobe right GM | cortical GM |
| Front.GM | Frontal lobe left GM | cortical GM |
| Pariet.GM | Parietal lobe right GM | cortical GM |
| Pariet.GM | Parietal lobe left GM | cortical GM |
| Caud | Caudate nucleus right | subcortical GM |
| Caud | Caudate nucleus left | subcortical GM |
| Thal.High | Thalamus right | subcortical GM |
| Thal.High | Thalamus left | subcortical GM |
| Subthal | Subthalamic nucleus right | subcortical GM |
| Subthal | Subthalamic nucleus left | subcortical GM |
| Lenti | Lentiform Nucleus right | subcortical GM |
| Lenti | Lentiform Nucleus left | subcortical GM |
| ATempMed.WM | Anterior temporal lobe | cortical GM |
| ATempMed.WM | Anterior temporal lobe | cortical GM |
| ATempLat.WM | Anterior temporal lobe | cortical GM |
| ATempLat.WM | Anterior temporal lobe | cortical GM |
| ParaAmbAnt.WM | Gyri parahippocampalis et ambiens anterior part left WM | WM |
| ParaAmbAnt.WM | Gyri parahippocampalis et ambiens anterior part right WM | WM |
| STempMid.WM | Superior temporal gyrus | WM |
| STempMid.WM | Superior temporal gyrus | WM |
| MITempAnt.WM | Medial and inferior temporal gyri anterior part left WM | WM |
| MITempAnt.WM | Medial and inferior temporal gyri anterior part right WM | WM |
| OccTempFusiAnt.WM | Lateral occipitotemporal gyrus | WM |

| | | |
|---|---|---|
| OccTempFusiAnt.WM | Lateral occipitotemporal gyrus | WM |
| Insula.WM | Insula right WM | WM |
| Insula.WM | Insula left WM | WM |
| Occ.WM | Occipital lobe right WM | WM |
| Occ.WM | Occipital lobe left WM | WM |
| ParaAmbPos.WM | Gyri parahippocampalis et ambiens posterior part right WM | WM |
| ParaAmbPos.WM | Gyri parahippocampalis et ambiens posterior part left WM | WM |
| OccTempFusiPos.WM | Lateral occipitotemporal gyrus | WM |
| OccTempFusiPos.WM | Lateral occipitotemporal gyrus | WM |
| MITempPos.WM | Medial and inferior temporal gyri posterior part right WM | WM |
| MITempPos.WM | Medial and inferior temporal gyri posterior part left WM | WM |
| STempPos.WM | Superior temporal gyrus | WM |
| STempPos.WM | Superior temporal gyrus | WM |
| CingAnt.WM | Cingulate gyrus | WM |
| CingAnt.WM | Cingulate gyrus | WM |
| CingPos.WM | Cingulate gyrus | WM |
| CingPos.WM | Cingulate gyrus | WM |
| Front.WM | Frontal lobe right WM | WM |
| Front.WM | Frontal lobe left WM | WM |
| Pariet.WM | Parietal lobe right WM | WM |
| Pariet.WM | Parietal lobe left WM | WM |
| Thal.Low | Thalamus right | subcortical GM |
| Thal.Low | Thalamus left | subcortical GM |

*GM = gray matter, WM = white matter.*

*Table S2 List of regional metrics selected by the baseline model based on individual metrics for the age prediction task.*

| |
|---|
| ***Volume:*** *Anterior temporal lobe, medial part right GM; Medial and inferior temporal gyri anterior part right GM; Anterior temporal lobe, medial part left WM; Anterior temporal lobe, lateral part left WM; Superior temporal gyrus, middle part right WM; Frontal lobe right WM; Parietal lobe left WM;* |
| ***FA:*** *Amygdala left; Anterior temporal lobe, medial part left GM; Gyri parahippocampalis et ambiens anterior part right GM; Superior temporal gyrus, middle part left GM; Superior temporal gyrus, middle part right GM; Cerebellum left; Cingulate gyrus, posterior part right GM; Cingulate gyrus, posterior part left GM; Parietal lobe left GM; Gyri parahippocampalis et ambiens posterior part left WM; Lateral occipitotemporal gyrus, gyrus fusiformis posterior part right WM; Cingulate gyrus, anterior part left WM;* |
| ***MD****: Medial and inferior temporal gyri posterior part right GM; Medial and inferior temporal gyri posterior part left GM; Superior temporal gyrus, posterior part right GM; Cingulate gyrus, posterior part right GM; Cingulate gyrus, posterior part left GM; Frontal lobe right GM; Caudate nucleus left; Thalamus left, high intensity part in T2; Subthalamic nucleus right; Subthalamic nucleus left ; Lentiform Nucleus left; Anterior temporal lobe, medial part right WM; Superior temporal gyrus, posterior part right WM; Superior temporal gyrus, posterior part left WM; Cingulate gyrus, posterior part left WM; Frontal lobe left WM; Parietal lobe right WM; Parietal lobe left WM;* |
| ***AD:*** *Hippocampus left; Hippocampus right; Amygdala left; Amygdala right; Gyri parahippocampalis et ambiens anterior part left GM; Gyri parahippocampalis et ambiens anterior part right GM; Superior temporal gyrus, middle part left GM; Lateral occipitotemporal gyrus, gyrus fusiformis anterior part right GM; Cerebellum left; Cerebellum right; Brainstem, spans the midline; Occipital lobe left GM; Gyri parahippocampalis et ambiens posterior part right GM; Superior temporal gyrus, posterior part right GM; Superior temporal gyrus, posterior part left GM; Anterior temporal lobe, medial part right WM; Anterior temporal lobe, lateral part left WM; Gyri parahippocampalis et ambiens anterior part left WM; Gyri parahippocampalis et ambiens anterior part right WM; Superior temporal gyrus, middle part left WM; Medial and inferior temporal gyri anterior part left WM;* |
| ***KURT:*** *Anterior temporal lobe, lateral part left GM; Anterior temporal lobe, lateral part right GM; Medial and inferior temporal gyri anterior part right GM; Lateral occipitotemporal gyrus, gyrus fusiformis anterior part left GM; Insula left GM; Medial and inferior temporal gyri posterior part right GM; Caudate nucleus right; Lentiform Nucleus right; Corpus Callosum; Lateral occipitotemporal gyrus, gyrus fusiformis anterior part right WM; Insula left WM; Medial and inferior temporal gyri posterior part right WM;* |
| ***ISO:*** *Amygdala left; Gyri parahippocampalis et ambiens anterior part left GM; Lateral occipitotemporal gyrus, gyrus fusiformis anterior part right GM; Gyri parahippocampalis et ambiens posterior part right GM; Gyri parahippocampalis et ambiens posterior part left GM; Superior temporal gyrus, posterior part left GM; Gyri parahippocampalis et ambiens posterior part right WM; Superior temporal gyrus, posterior part left WM;* |
| ***ODIp:*** *Anterior temporal lobe, medial part right GM; Insula left GM; Medial and inferior temporal gyri posterior part right GM; Medial and inferior temporal gyri posterior part left GM;* |

*Table S3 List of regional metrics selected by the baseline model based on individual metrics for the classification task.*

| |
|---|
| ***Volume:*** *Anterior temporal lobe, lateral part right GM; Superior temporal gyrus, middle part right GM; Medial and inferior temporal gyri anterior part right GM; Lateral occipitotemporal gyrus, gyrus fusiformis anterior part left GM; Lateral occipitotemporal gyrus, gyrus fusiformis anterior part right GM; Brainstem, spans the midline; Insula right GM; Caudate nucleus left; Lateral occipitotemporal gyrus, gyrus fusiformis anterior part left WM; Occipital lobe left WM; Medial and inferior temporal gyri posterior part right WM; Medial and inferior temporal gyri posterior part left WM; Frontal lobe right WM; Frontal lobe left WM; Parietal lobe left WM;* |
| ***T1/T2:*** *Hippocampus left; Superior temporal gyrus, middle part left GM; Superior temporal gyrus, middle part right GM; Medial and inferior temporal gyri anterior part left GM; Lateral occipitotemporal gyrus, gyrus fusiformis anterior part left GM; Lateral occipitotemporal gyrus, gyrus fusiformis anterior part right GM; Cingulate gyrus, anterior part right GM; Parietal lobe left GM; Caudate nucleus right; Thalamus left, high intensity part in T2; Anterior temporal lobe, lateral part left WM; Gyri parahippocampalis et ambiens posterior part left WM; Medial and inferior temporal gyri posterior part right WM;* |
| ***FA:*** *Gyri parahippocampalis et ambiens posterior part right GM; Gyri parahippocampalis et ambiens posterior part left GM; Medial and inferior temporal gyri posterior part right GM; Frontal lobe right GM; Parietal lobe right GM; Subthalamic nucleus left ; Insula left WM;* |
| ***MD:*** *Amygdala right; Lateral occipitotemporal gyrus, gyrus fusiformis anterior part left GM; Cingulate gyrus, posterior part right GM; Anterior temporal lobe, medial part left WM; Anterior temporal lobe, lateral part right WM; Occipital lobe left WM; Lateral occipitotemporal gyrus, gyrus fusiformis posterior part left WM; Medial and inferior temporal gyri posterior part left WM; Superior temporal gyrus, posterior part right WM; Superior temporal gyrus, posterior part left WM; Cingulate gyrus, posterior part right WM; Parietal lobe left WM;* |
| ***AD:*** *Amygdala right; Cerebellum right; Gyri parahippocampalis et ambiens posterior part right WM; Cingulate gyrus, posterior part left WM;* |
| ***RD:*** *Occipital lobe right GM; Lateral occipitotemporal gyrus, gyrus fusiformis posterior part left GM; Lentiform Nucleus left; Lateral occipitotemporal gyrus, gyrus fusiformis anterior part left WM; Lateral occipitotemporal gyrus, gyrus fusiformis posterior part left WM; Medial and inferior temporal gyri posterior part right WM; Superior temporal gyrus, posterior part right WM; Thalamus left, low intensity part in T2;* |
| ***KURT:*** *Gyri parahippocampalis et ambiens anterior part right GM; Superior temporal gyrus, middle part left GM; Superior temporal gyrus, middle part right GM; Cerebellum left; Insula left GM; Occipital lobe right GM; Occipital lobe left GM; Medial and inferior temporal gyri posterior part right GM; Medial and inferior temporal gyri anterior part left WM; Frontal lobe left WM; Thalamus right, low intensity part in T2;* |
| ***ICVF:*** *Hippocampus right; Anterior temporal lobe, medial part right GM; Anterior temporal lobe, lateral part left GM; Lateral occipitotemporal gyrus, gyrus fusiformis anterior part left GM; Lateral occipitotemporal gyrus, gyrus fusiformis anterior part right GM; Brainstem, spans the midline; Caudate nucleus left; Gyri parahippocampalis et ambiens anterior part left WM; Occipital lobe right WM;* |
| ***ISO:*** *Superior temporal gyrus, middle part left GM; Superior temporal gyrus, middle part right GM; Lentiform Nucleus left; Superior temporal gyrus, middle part left WM; Lateral occipitotemporal gyrus, gyrus fusiformis anterior part left WM; Medial and inferior temporal gyri posterior part left WM; Superior temporal gyrus, posterior part right WM; Cingulate gyrus, anterior part right WM; Thalamus right, low intensity part in T2; Thalamus left, low intensity part in T2;* |
| ***ODItot:*** *Superior temporal gyrus, middle part left GM; Insula right GM; Occipital lobe right GM; Cingulate gyrus, posterior part left GM; Caudate nucleus right; Caudate nucleus left; Thalamus left, high intensity part in T2; Medial and inferior temporal gyri anterior part right WM; Gyri parahippocampalis et ambiens posterior part right WM; Gyri parahippocampalis et ambiens posterior part left WM; Frontal lobe left WM; Parietal lobe left WM;* |
| ***ODIp:*** *Amygdala right; Anterior temporal lobe, medial part right GM; Anterior temporal lobe, lateral part right GM; Gyri parahippocampalis et ambiens anterior part left GM; Cerebellum right; Occipital lobe left GM; Gyri parahippocampalis et ambiens posterior part left GM; Lateral occipitotemporal gyrus, gyrus fusiformis posterior part right GM; Medial and inferior temporal gyri posterior part left GM; Frontal lobe right GM; Caudate nucleus right; Anterior temporal lobe, lateral part right WM; Insula left WM; Lateral occipitotemporal gyrus, gyrus fusiformis posterior part left WM; Parietal lobe right WM;* |
